# Apical cell expansion maintained by Dusky-like establishes a scaffold for corneal lens morphogenesis

**DOI:** 10.1101/2024.01.17.575959

**Authors:** Neha Ghosh, Jessica E. Treisman

## Abstract

The biconvex shape of the *Drosophila* corneal lens, which enables it to focus light onto the retina, arises by organized assembly of chitin and other apical extracellular matrix components. We show here that the Zona Pellucida domain-containing protein Dusky-like is essential for normal corneal lens morphogenesis. Dusky-like transiently localizes to the expanded apical surfaces of the corneal lens-secreting cells, and in its absence, these cells undergo apical constriction and apicobasal contraction. Dusky-like also controls the arrangement of two other Zona Pellucida-domain proteins, Dumpy and Piopio, external to the developing corneal lens. Loss of either *dusky-like* or *dumpy* delays chitin accumulation and disrupts the outer surface of the corneal lens. Artificially inducing apical constriction with constitutively active Myosin light chain kinase is sufficient to similarly alter chitin deposition and corneal lens morphology. These results demonstrate the importance of cell shape for the morphogenesis of overlying apical extracellular matrix structures.

## Introduction

The extracellular matrix (ECM) is a complex array of proteins and polysaccharides that provides structural and biochemical support to tissues. Mutations affecting the ECM can cause numerous human genetic disorders including cancer, muscular dystrophy, hematuria, and retinopathy ^1–3^. Basement membrane ECM, which forms a sheet underlying epithelial cells, is made up of conserved proteins such as collagen, nidogen, perlecan, and laminins. It can mediate adhesion between tissue layers, insulate cells from the extracellular fluid, transmit mechanical forces, influence the distribution of signaling molecules, and act as a substrate for cell migration ^4,5^. Less is known about the apical ECM (aECM), which has diverse forms and functions.

External aECMs such as invertebrate cuticles can act as permeability barriers and protect against desiccation, and the shapes of tubular organs such as the *Drosophila* trachea and *C. elegans* excretory system depend on luminal scaffolds composed of aECM ^6,7^. In vertebrates, aECM forms structures such as the tectorial membrane of the inner ear, the vascular glycocalyx, pulmonary surfactant, and the mucin-rich lining of the gut ^8–13^. The morphogenesis of aECM assemblies is not fully understood, although in a few cases it has been linked to cytoskeletal organization in the underlying cells ^14,15^.

One of the most striking structures composed of aECM is the *Drosophila* corneal lens, which consists of fibers of the polysaccharide chitin and associated proteins arranged to form a precisely curved biconvex shape ^16,17^. The corneal lens of each ommatidium of the compound eye is secreted during the second half of pupal development by the underlying non-neuronal cells^18^. Major corneal lens constituents such as Crystallin are produced by the central cone and primary pigment cells, while additional components are derived from the secondary and tertiary pigment cells attached to the corneal lens periphery ^16,19^. Although it is not known how these cells specify the shape of the corneal lens, we have previously shown that the transcription factor Blimp-1 is essential to generate its outer curvature ^20^. Many Blimp-1 target genes encode aECM components, including members of the most conserved family of aECM proteins, which are characterized by a Zona Pellucida (ZP) domain ^20,21^.

The ZP domain was initially identified in constituents of the extracellular coat surrounding the mammalian oocyte ^22,23^, but has since been found in proteins implicated in many developmental processes ^21,24^. Mutations affecting human ZP-domain proteins are associated with disease conditions that include infertility, deafness, vascular disorders, inflammatory bowel disease, kidney disease, and cancer ^22,25–30^. The 260 amino acid ZP domain can mediate homomeric or heteromeric polymerization into filaments; it consists of N- and C-terminal subdomains with structures related to immunoglobulin domains that are stabilized by conserved disulfide bonds ^31–33^. Some ZP-domain proteins are tethered to the plasma membrane by a transmembrane domain or GPI linkage, while others undergo proteolytic cleavage and are secreted into the extracellular space ^34–37^.

ZP-domain proteins have been found to play a crucial role in shaping the aECM and attaching it to the apical plasma membrane ^7,21^. For instance, the α-tectorin ZP-domain protein is essential for attachment of the tectorial membrane to the cochlear epithelium ^38^. Filaments of the giant *Drosophila* ZP-domain protein Dumpy (Dpy) anchor pupal appendages and tendon cells to the external cuticle ^39–41^, and the dendrites of sensory organs also require ZP-domain proteins to connect them to the cuticle ^37,42^. The lumen of tubular structures in *Drosophila, C. elegans*, and the mammalian kidney is organized by ZP-domain proteins ^29,43–45^. Interestingly, mutations affecting eight *Drosophila* ZP-domain proteins each have distinct effects on the morphology of embryonic denticles, and each protein occupies a specific spatial location in the denticle ^46^, indicating that ZP-domain proteins are specialized to assemble aECM into diverse shapes.

Here, we show that the ZP-domain protein Dusky-like (Dyl) ^46–48^ is essential for normal morphogenesis of the *Drosophila* corneal lens. During pupal development, *dyl* mutant ommatidia undergo abnormal apical constriction and apicobasal contraction accompanied by changes in the distribution of cytoskeletal proteins. Loss of *dyl* also results in changes in the organization of other ZP-domain proteins such as Dpy and Piopio (Pio) ^36,43,49^, and a delay in the accumulation of chitin. Interestingly, many of these changes can be phenocopied by activating myosin to artificially induce constriction of the corneal lens-secreting cells. These observations suggest that Dyl acts as a mechanical anchor that transiently attaches the cone and primary pigment cells to the aECM to maintain their expanded apical surface area, enabling the generation of a ZP-domain protein scaffold that shapes the curved corneal lens surface.

## Results

### Dyl is necessary for normal corneal lens morphology

The *Drosophila* corneal lens is a biconvex aECM structure that focuses light onto the underlying photoreceptors ^17^. The transcription factor Blimp-1 is necessary for normal corneal lens curvature, and its target genes include several that encode ZP-domain proteins ^20^. ZP-domain proteins have been reported to connect aECM to the plasma membrane of epithelial cells, and mutations affecting these proteins can alter the shape of cuticular structures such as embryonic denticles ^46^ and bristles ^47,48^. We therefore used existing mutants and RNAi lines to test whether any ZP-domain targets of Blimp-1 were necessary for the normal shape of the corneal lens.

We observed a dramatic effect on the external appearance of the adult eye in clones mutant for one such gene, *dyl*. Scanning electron micrographs showed that in ommatidia homozygous for *dyl^Δ1^*^42^, a deletion of the entire gene ^46^, the corneal lenses lacked external curvature and had gaps in their outer surfaces (Fig. 1A). The same flat shape and abnormal surface structure were visible in transmission electron micrographs (Fig. S1A, B). Cryosections of eyes stained for chitin with Calcofluor White revealed that the flatter external surfaces of the *dyl* mutant corneal lenses were accompanied by more deeply curved internal surfaces than those of the adjacent wild-type biconvex corneal lenses (Fig. 1C). Quantification of the outer and inner angles between adjacent corneal lenses (Fig. 1E) confirmed that the outer angle was significantly increased and the inner angle significantly decreased in *dyl* mutant regions (Fig. 1F, G). The changes in corneal lens shape observed in *dyl^Δ1^*^42^ mutants, which included increased height and reduced width, were rescued by expression of a *UAS-dyl* transgene ^46^ in all retinal cells with *lGMR-GAL4* ^50^, confirming that the defects were due to loss of *dyl* (Fig. 1B, D, F, G, Fig. S1E, F). In transmission electron micrographs, we observed that the deeper inner curvature of the corneal lens correlated with an increase in its area of contact with the peripheral secondary and tertiary pigment cells (Fig. S1A-D, G). Taken together, our results show that the ZP-domain protein Dyl is necessary for the normal biconvex shape and external morphology of the corneal lens.

**Figure 1.**
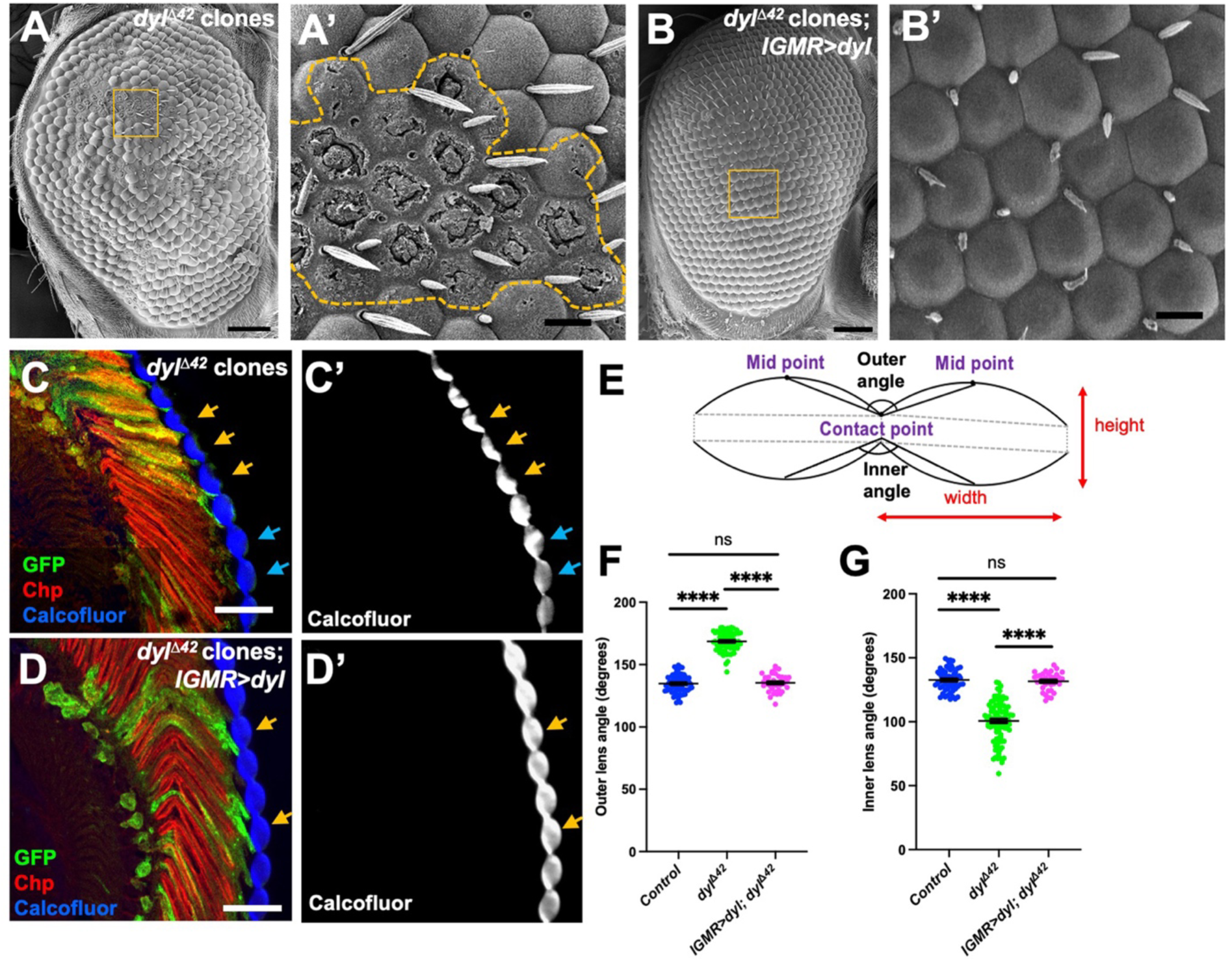
dyl is necessary for normal corneal lens curvature. (**A, B**) Scanning electron micrographs of adult eyes. (**A**) in *dyl**^Δ^***^42^ mutant clones, the corneal lens surfaces are flat rather than convex, and many appear to be missing parts of the surface layer. (**B**) *dyl**^Δ^***^42^ mutant clones expressing a *UAS-dyl* transgene with *lGMR-GAL4* cannot be distinguished from wild-type eye tissue. The regions in the yellow boxes are enlarged in (**A’, B’**), and the boundaries of the *dyl* mutant clone are marked with a yellow dotted line in (**A’**). (**C, D**) Horizontal frozen sections of adult eyes in which *dyl**^Δ^***^42^ mutant clones (**C**) or *dyl**^Δ^***^42^ mutant clones expressing *UAS-dyl* with *lGMR-GAL4* (**D**) are positively marked with GFP (green). The corneal lenses are stained with Calcofluor White (**C’, D’,** blue in **C, D**) and photoreceptors are marked with anti-Chaoptin (Chp, red). *dyl* mutant ommatidia have externally flat corneal lenses (yellow arrows, **C**) while adjacent wild-type ommatidia have biconvex corneal lenses (blue arrows, **C**). Rescue with *UAS-dyl* restores the normal corneal lens shape (yellow arrows, **D**). (**E**) Schematic illustration showing how the outer and inner angles between adjacent corneal lenses and the height and width of a corneal lens were defined. (**F, G**) Graphs showing the outer (**F**) and inner (**G**) angles between adjacent corneal lenses in adult eye sections for wild type control regions, *dyl**^Δ^***^42^ mutant clones, and *dyl**^Δ^***^42^ mutant clones in which *UAS-dyl* is driven with *lGMR-GAL4*. For both graphs, error bars show mean ± SEM. n values are given as number of ommatidia/number of retinas for this and subsequent graphs. Wild type n=70/20, *dyl* mutant n=103/20, *lGMR>dyl*; *dyl* mutant n=37/6. ****p < 0.0001, unpaired two-tailed t test with Welch’s correction. Scale bars: 200 μm (**A**, **B**), 2.5 μm (**A’**, **B’**), 20 µm (**C, D**).

### Loss of *dyl* causes apical constriction and apico-basal contraction of ommatidia

In order to understand how loss of *dyl* affects corneal lens shape, we traced the mutant phenotype earlier in development. Corneal lens secretion initiates at 50 h APF and continues through the late pupal stages ^18^. Major components of the corneal lens such as Crystallin and Retinin are produced by the underlying cone and primary pigment cells, and additional constituents are secreted by the secondary and tertiary pigment cells (also called lattice cells, Fig. 2G) ^16^. To determine the site of Dyl expression in the retina, we used an antibody against Dyl ^46^, and identified specific labeling by its absence in *dyl* mutant clones (Fig. 2A, B, Fig. S2A, B). We observed that Dyl was present on the apical surface of the cone and primary pigment cells at 48 h APF (Fig. 2A) and 50 h APF (Fig. S2A). Dyl protein levels were reduced at 52 h APF (Fig. S2B) and became very low at 54 h APF (Fig. 2B), consistent with the transient 48 h APF peak of *dyl* mRNA reported by modENCODE ^51^.

**Figure 2.**
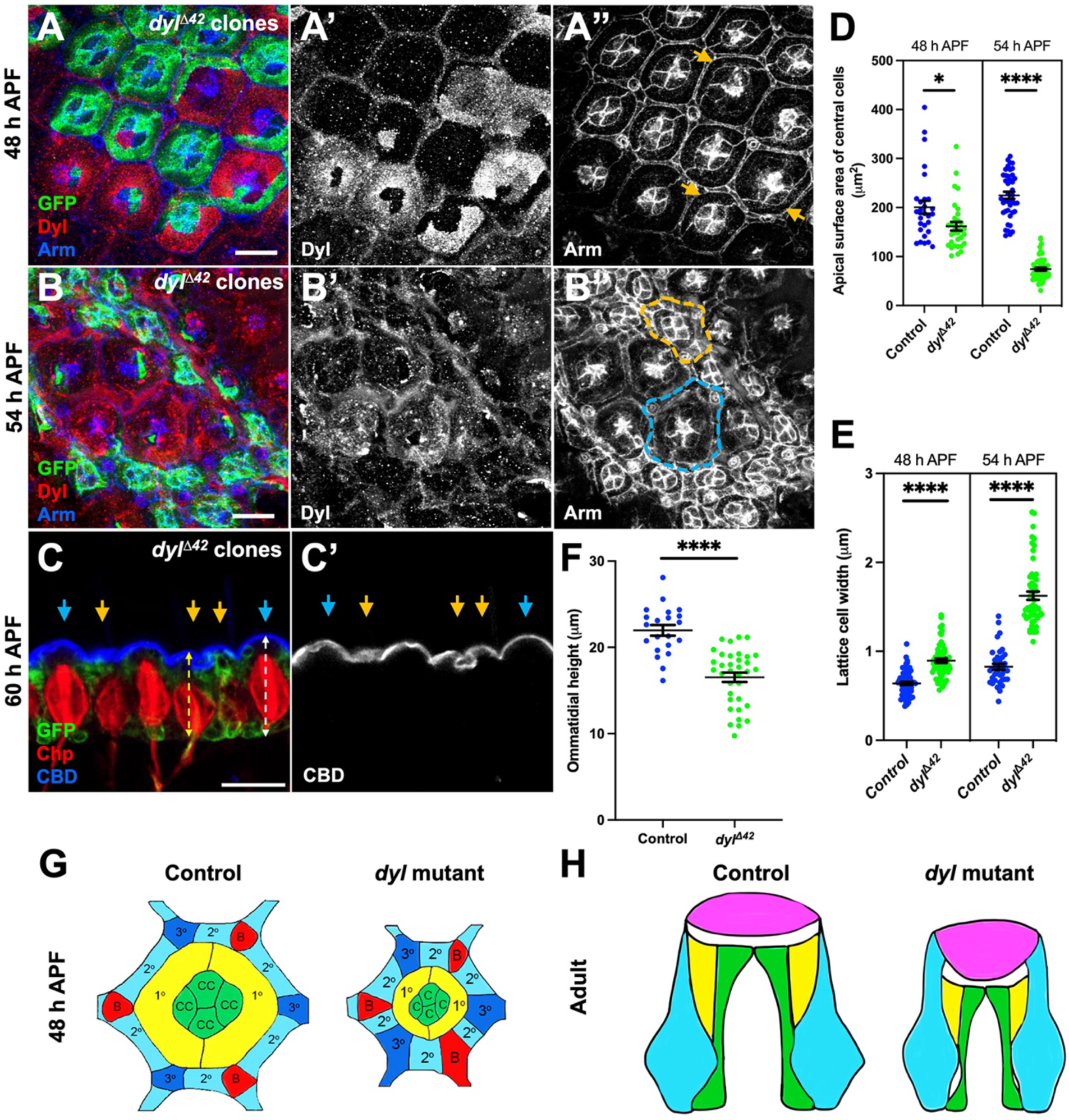
Loss of *dyl* causes apical constriction and apico-basal contraction of ommatidia. (**A, B**) Retinas containing *dyl**^Δ^***^42^ mutant clones marked with GFP (green) and stained with anti-Dyl (**A’**, **B’**, red in **A**, **B**) and anti-Armadillo (Arm) to mark apical junctions (**A’’**, **B’’**, blue in **A**, **B**) at (**A**) 48 h APF and (**B**) 54 h APF. The *dyl**^Δ^***^42^ mutant clones show loss of Dyl staining in cone and primary pigment cells (there is non-specific staining of the lattice cells), and a marked reduction in their apical surface area (yellow arrows, **A”**; yellow outline, **B’’**) as well as apicobasal contraction; the apical plane of the *dyl* mutant ommatidium outlined in yellow is in the same confocal section as a more basal plane of the wild-type ommatidium outlined in blue (**B”**). (**C**) Horizontal cryosection of a 60h APF retina showing apicobasal contraction in *dyl**^Δ^***^42^ mutant clones marked with GFP (green, yellow arrows) in comparison to wild type ommatidia (blue arrows). An Alexa647-labeled chitin binding domain (CBD, **C’,** blue in **C**) marks the corneal lenses, which are flattened and distorted in *dyl* mutant clones, and photoreceptor rhabdomeres are stained with anti-Chp (red). Dotted lines show how ommatidial height was measured. Scale bars: 10 µm (**A, B**), 20 µm (**C**). (**D, E**) Graphs depicting (**D**) apical surface area of the central cone and primary pigment cells and (**E**) lattice cell width in *dyl**^Δ^***^42^ mutant clones and control wild-type ommatidia in the same retinas, at 48 h APF (n=28/18 for control and 32/18 for *dyl* in **D**, n=69/7 for control and 61/7 for *dyl* in **E**) and 54 h APF (n=44/27 for control and 52/12 for *dyl* in **D**, n=37/7 for control and 57/7 for *dyl* in **E**). (**F**) graph showing ommatidial height in *dyl**^Δ^***^42^ mutant clones and control wild-type ommatidia at 60 h APF (n=21/5 for control and 35/5 for *dyl*). For all graphs, error bars show mean ± SEM. ****p < 0.0001, *p = 0.0221, unpaired two-tailed t test with Welch’s correction. (**G**) Schematic representations of ommatidia at 54 h APF. Cone cells (CC, green) and primary pigment cells (1°, yellow), together known as the central cells, are surrounded by a lattice of secondary pigment cells (2°, cyan), tertiary pigment cells (3°, indigo), and bristles (B, red). In controls, we hypothesize that Dyl attaches the central cells to the aECM to maintain their apical surfaces in an expanded state. In *dyl* mutants, the central cells would lose their apical attachments and contract. (**H**) Schematic of horizontal views of control and *dyl* mutant adult ommatidia. The corneal lens (pink) may maintain its attachment to the lattice cells as the ommatidium contracts basally, flattening its external curvature and deepening its internal curvature.

*dyl* mutant ommatidia showed a striking apical constriction starting at 48 h APF (Fig. 2A) that became quite pronounced at 54 h APF (Fig. 2B, Fig. S2F). The change was driven by a large decrease in the apical surface area of the cone and primary pigment cells (central cells) (Fig. 2D); in contrast, the apical width of the lattice cells expanded (Fig. 2E). The apical constriction of *dyl* mutant ommatidia was accompanied by apical-basal contraction, placing their apical surfaces in a deeper plane than the wild-type ommatidia at 54 h APF (Fig. 2B). This change could also be visualized in cryosections of 60 h APF pupal retinas, in which *dyl^Δ^*^42^ ommatidia were shorter than their wild-type neighbors (Fig. 2C, F). In these sections, the wild-type corneal lenses labeled with a fluorescent chitin-binding domain (CBD) had convex outer surfaces, whereas corneal lenses corresponding to *dyl* clones were flat or convoluted (Fig. 2C). Expression of the wild-type UAS-*dyl* transgene rescued both the apical constriction and apicobasal contraction displayed by *dyl^Δ^*^42^ mutants (Fig. S2C-F). These data suggest that Dyl acts at the apical surfaces of cone and primary pigment cells to maintain their apical expansion (Fig. 2G). The apicobasal contraction of *dyl* mutant ommatidia correlates with a change in the shape of the developing corneal lens (Fig. 2H).

### Loss of *dyl* causes cytoskeletal reorganization

We next investigated how the organization of the cytoskeleton was affected in *dyl* mutant clones. Using the actin-binding domain of the ERM protein Moesin (Moe) tagged with mCherry ^52^ to label actin filaments, we found that at 60 h APF more Moe::mcherry accumulated at the apical surface of *dyl* mutant ommatidia than in neighboring wild-type ommatidia, implying that the actin cytoskeleton is condensed in *dyl* mutant cells (Fig. 3A). Actomyosin contraction can be induced by phosphorylation of the non-muscle myosin II regulatory light chain, known in *Drosophila* as Spaghetti squash (Sqh) ^53^. We stained *dyl^11^*^42^ clones in the pupal retina with an antibody against phospho-Sqh (pSqh) ^54^ and found it to be enriched in lattice cells at 50 h APF in both wild-type and *dyl* mutant regions (Fig. S3A). At 54 h APF pSqh was present in discrete foci at the apical surfaces of wild-type cone and primary pigment cells, whereas in *dyl^Δ^*^42^ clones, pSqh was more uniformly distributed (Fig. 3B). Loss of *dyl* had a similar effect on the distribution of beta-heavy spectrin (β_H_-spectrin) (Fig. 3C, S3B), a component of the spectrin-based membrane cytoskeleton encoded by the gene *karst* (*kst*) that has been reported to control apical cell surface area ^55^. Similar foci of phospho-Sqh and β_H_-spectrin have been observed in cells undergoing pulsatile apical constriction and ratcheting, in the embryonic mesoderm and other contexts ^56,57^. Myosin foci are also associated with fluctuations in the area of cone and primary pigment cells as they expand earlier in pupal development ^58^. The lack of such foci in *dyl* mutant cells may indicate that their constriction is a less organized process.

**Figure 3.**
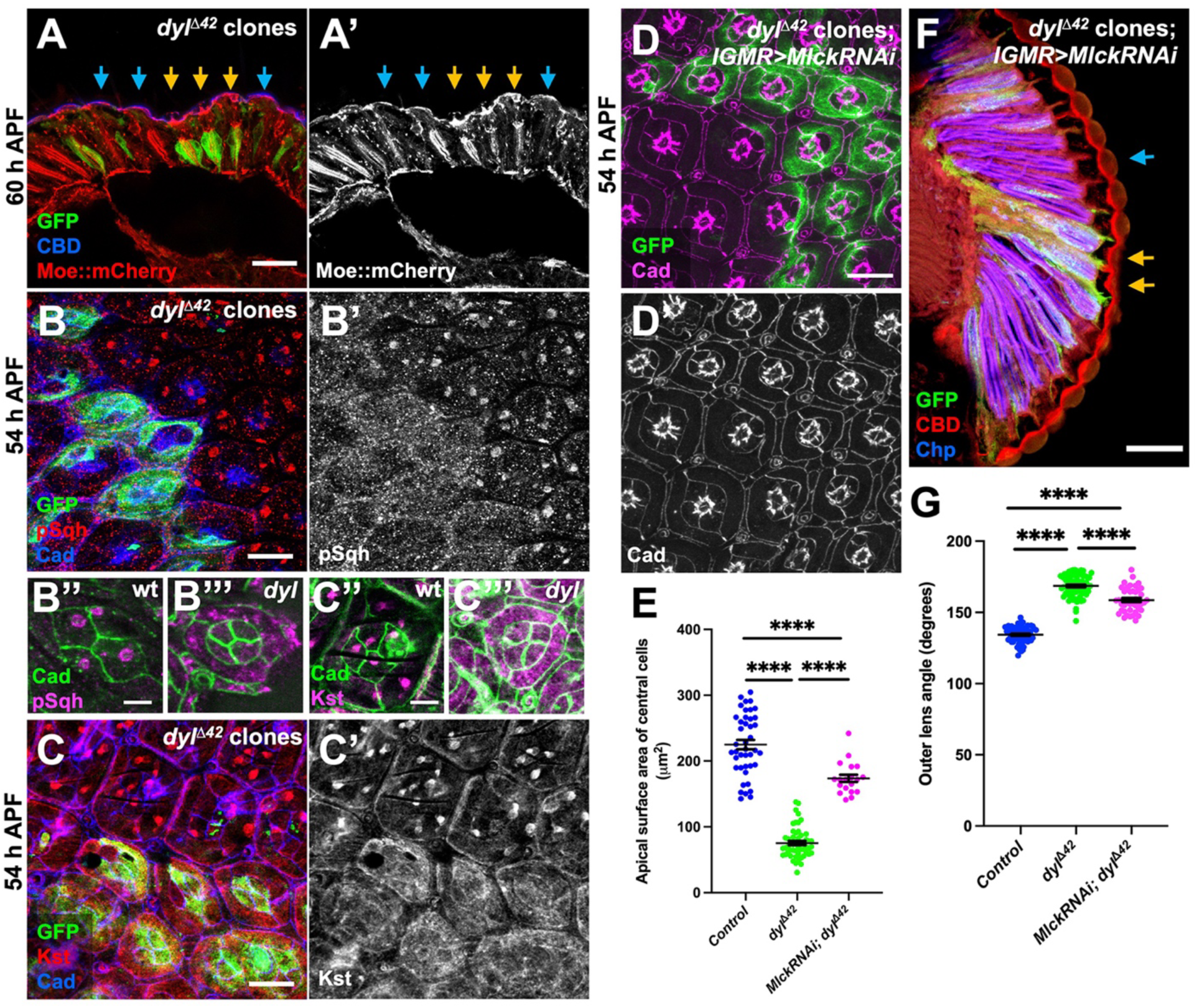
Loss of *dyl* alters the distribution of cytoskeletal proteins. (**A**) Cryosection of a 60 h APF retina showing Moe::mCherry (red) accumulation at the apical surface in *dyl**^Δ^***^42^ mutant clones (yellow arrows) marked with GFP (green). Blue arrows mark wild type ommatidia. CBD (blue) labels the corneal lens. (**B, C**) 54 h APF retinas containing *dyl**^Δ^***^42^ mutant clones marked with GFP (green), stained with anti-Ecad and anti-Ncad (blue), anti-pSqh (**B’**, red in **B**), or anti-β_H_-spectrin (**C’**, red in **C**). In wild-type ommatidia at this stage pSqh and β_H_-spectrin form foci, but in *dyl**^Δ^***^42^ mutant clones they are diffusely localized. Confocal sections showing the apical surfaces of individual wild-type (**B’’, C’’**) or *dyl* mutant (**B’’’, C’’’**) ommatidia labeled with anti-Ecad and anti-Ncad (green) and anti-pSqh (magenta, **B’’, B’’’**) or anti-βH-spectrin (magenta, **C’’, C’’’**) show that the differences are not due to the more basal position of *dyl* mutant ommatidia. (**D**) A 54 h APF retina with *dyl**^Δ^***^42^ mutant clones expressing *UAS-Mlck RNAi* with *lGMR-GAL4*, marked with GFP (green). Apical cell junctions are stained with anti-Ecad and anti-Ncad (**D’**, magenta in **D**). (**E**) Graph showing the apical surface area of central cells in *dyl**^Δ^***^42^ mutant clones and internal wild type control ommatidia (data from Fig. 2D), and in *dyl**^Δ^***^42^ mutant clones expressing *Mlck RNAi* (n=19/8) at 54 h APF. Knocking down *Mlck* partially rescues the apical constriction phenotype. (**F**) Horizontal cryosection of an adult eye with *dyl**^Δ^***^42^ mutant clones expressing *Mlck RNAi* labeled with GFP (green, yellow arrows), stained with anti-Chp (blue) and CBD (red). Scale bars: 10 µm (**A-D**), 5 μm (**B’’, B’’’, C’’, C’’’**), 20 μm (**F**). (**G**) Graph showing the outer angle between adjacent corneal lenses in control, *dyl**^Δ^***^42^ mutant clones (data from Fig. 1F), and *dyl**^Δ^***^42^ mutant clones expressing *Mlck RNAi* (n=41/8). Knocking down *Mlck* is not sufficient to restore normal corneal lens curvature in *dyl**^Δ^***^42^ mutant clones. Error bars show mean +/-SEM. ****p < 0.0001, unpaired two-tailed t test with Welch’s correction.

### Artificially induced apical constriction alters corneal lens shape

We attempted to test whether apical constriction was necessary for the effect of *dyl* on corneal lens shape by knocking down *Myosin light chain kinase* (*Mlck)*, which encodes a kinase that phosphorylates Sqh to induce actomyosin contraction ^59^, in *dyl^Δ^*^42^ clones. A significant change in corneal lens shape was still observed, but because the apical constriction was not fully rescued, we could not determine whether *dyl* has an effect on corneal lens shape that is independent of apical constriction (Fig. 3D-G). Knocking down *kst* also partially rescued the apical constriction of *dyl* mutant ommatidia, but did not restore normal corneal lens curvature (Fig. S3C-F). These knockdowns may have been incomplete, or the constriction may be partially independent of Mlck and β_H_-spectrin activity. If Dyl-mediated attachments to the apical ECM exert tension on the apical surfaces of the cone and primary pigment cells to maintain their expansion, loss of these attachments could result in passive constriction of the apical surfaces.

We next tested whether apical constriction was sufficient to explain the effect of *dyl* on corneal lens shape. We artificially induced apical constriction by expressing a constitutively active form of Mlck (*UAS-Mlck^CT^*) ^60^ with *lGMR-GAL4* in clones in the retina. We confirmed that *Mlck^CT^*-expressing clones exhibited strong apical constriction in the mid-pupal retina (Fig. 4A). At the adult stage, the overlying corneal lenses were externally flat with deep and distorted internal curvature (Fig. 4B-D). Apical constriction in *dyl^Δ^*^42^ clones is limited to the cone and primary pigment cells (Fig. 2D, E). Expressing *UAS-Mlck^CT^*specifically in cone and primary pigment cells with *sparkling-GAL4* (*spa-GAL4*) ^61^ resulted in a change in pSqh distribution and corneal lens shape similar to *dyl* mutant clones, indicating that constriction of these cells is sufficient to alter corneal lens curvature (Fig. 4E-G).

**Figure 4.**
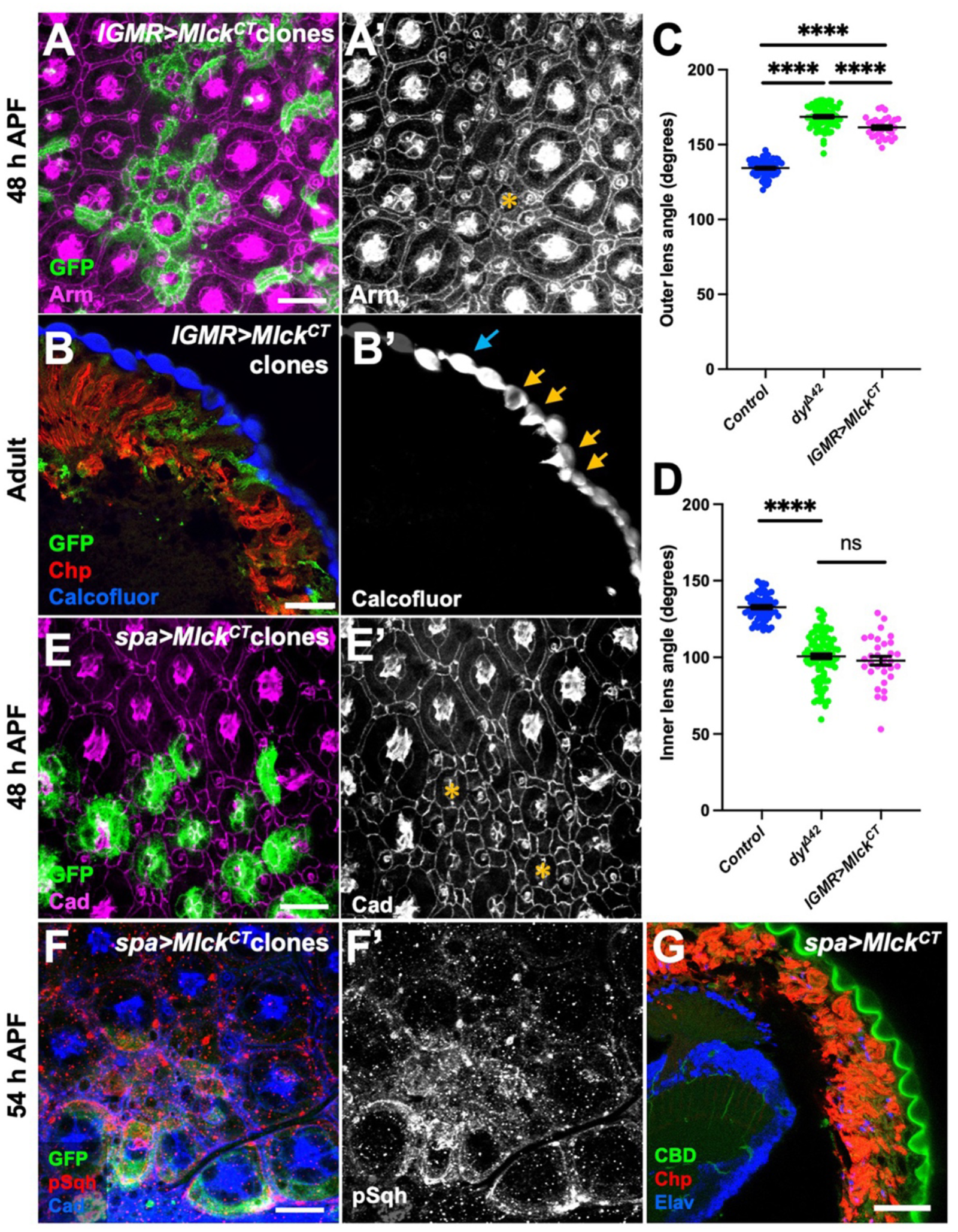
Apical constriction of central cells is sufficient to alter corneal lens shape. (**A, B**) Expression of a constitutively active form of *Mlck* (UAS-*Mlck^CT^*) in clones marked with GFP (green) in a 48 h APF pupal retina (**A**) and adult eye section (**B**) stained for Arm (**A’**, magenta in **A**), Chp (red in **B**) and Calcofluor White (B’, blue in B). UAS-*Mlck^CT^* clones display strong apical constriction of central cells (yellow asterisk, **A’**), and in the adult eye, they show distorted corneal lenses with flatter external surfaces and deeper internal curvature (yellow arrows in **B’**) than controls (blue arrow). (**C, D**) Graphs showing (**C**) outer angle and (**D**) inner angle between adjacent corneal lenses in control, *dyl**^Δ^***^42^ mutant clones (data from Fig. 1F, G), and *Mlck^CT^* overexpressing clones (n=31/5). Error bars show mean +/-SEM. ****p < 0.0001, ns p=0.369, unpaired two-tailed t test with Welch’s correction. (**E, F**) UAS-*Mlck^CT^* is expressed only in cone and primary pigment cells with *spa-GAL4* in clones marked with GFP (green) at 48 h APF (**E**) or 54 h APF (**F**). Retinas are stained with anti-E-Cad and anti-N-Cad (**E’**, magenta in **E,** blue in **F**) and anti-pSqh (F’, red in F). The clones show apical constriction of the central cells and expansion of the lattice cells (asterisks in **E’**) and accumulation of disorganized pSqh. (**G**) Horizontal cryosection of an adult eye in which *spa-GAL4* drives *UAS-MLCK^CT^* in all ommatidia, stained for Chp (red), CBD (green) and the neuronal nuclear marker Elav (blue). CBD staining shows external flattening and deep internal curvature of the corneal lenses. Scale bars: 10 µm (**A, E, F**), 20 µm (**B, G**).

### Dyl affects the organization of other ZP-domain proteins

The ZP domain is known to mediate homodimerization or heterodimerization, allowing ZP-domain proteins to form extended filaments. Previous studies have shown that the ZP-domain protein Dpy ^49^ interacts with another ZP-domain protein, Piopio (Pio), in the epidermis and the lumen of the tracheal system ^36,43^. Additionally, the ZP-domain protein Quasimodo (Qsm) modifies the strength of the Dpy filaments that connect tendon cells to the external pupal cuticle ^40^. Since Dyl is only transiently expressed in the developing pupal retina, we wondered whether it might interact with other ZP-domain proteins to maintain corneal lens structure. To test this hypothesis, we examined the effect of loss of *dyl* on the organization of Dpy and Pio, which we found to colocalize in the pupal retina (Fig. S5A). At 50 h APF, Dpy and Pio were uniformly localized above the cone and primary pigment cells in both wild type and *dyl^Δ^*^42^ ommatidia (Fig. 5A, Fig. S5B). By 54 h APF, Dpy and Pio were organized into a variety of linear structures above the central region of wild-type ommatidia, but maintained their uniform distribution in *dyl^Δ^*^42^ clones (Fig. 5B, G). Loss of *dyl* had a similar effect on another ZP-domain protein, Trynity (Tyn) ^46,62^ (Fig. S5C). These observations suggest that other ZP-domain proteins depend on Dyl for their normal organization.

**Figure 5.**
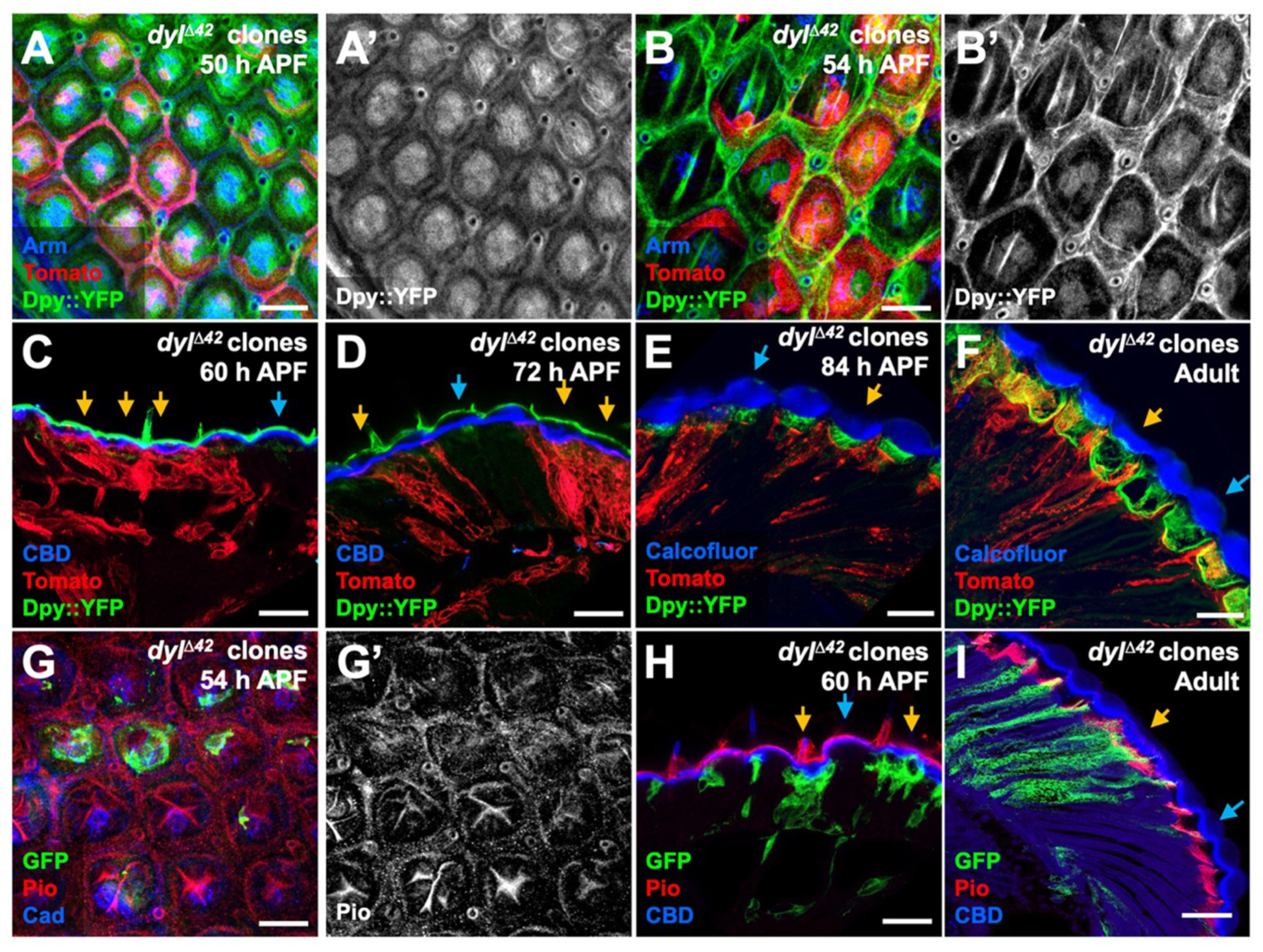
Loss of *dyl* affects the organization of Dpy and Pio at mid-pupal stages. (**A, B**) Pupal retinas at 50 h APF (**A**) and 54 h APF (**B**) with *dyl*^1142^ clones marked by myristoylated Tomato (red), stained for Dpy-YFP (**A’**, **B’**, green in **A**, **B**) and Arm (blue). Dpy-YFP takes on a linear organization that becomes more pronounced at later stages in wild-type but not *dyl* mutant ommatidia. **(C-F)** Horizontal sections of retinas containing *dyl*^1142^ clones marked by myristoylated Tomato (red) at 60 h APF (**C**), 72 h APF (**D**), 84 h APF (**E**) and adult stage (**F**), stained for Dpy-YFP (green) and CBD (blue, **C**, **D**) or Calcofluor White (blue, **E**, **F**). From 60-72 h APF Dpy-YFP is present external to the chitin layer of the corneal lenses and is less convex in *dyl* mutant clones (yellow arrows) than wild-type (blue arrows), but at 84 h APF it is lost from the external surface and begins to be secreted into the pseudocone under the corneal lens, where it remains in the adult. (**G-I**) *dyl**^Δ^***^42^ mutant clones marked with GFP (green) in a 54 h APF retina **(G)**, and in horizontal cryosections of 60 h APF **(H)** and adult **(I)** retinas, stained for Pio (**G’**, red in **G-I**), Ecad and Ncad (blue in **G**), or CBD (blue in **H**, **I**). Pio localizes in a similar pattern to Dpy-YFP in pupal retinas, but is restricted to the basal part of the pseudocone in adults. Dpy and Pio are also visible in mechanosensory bristles in (**C, D, H**). Scale bars: 10 µm (**A, B, G**), 20 µm (**C-F, H-I**).

Tyn, like Dyl, was only detected at mid-pupal stages, but Dpy and Pio were maintained into adulthood. At 60 h and 72 h APF, we found that Dpy and Pio were apical to chitin in the developing corneal lens, and had a less convex shape in *dyl^Δ^*^42^ clones than in wild-type ommatidia (Fig. 5C, D, H). However, when pseudocone secretion begins at 84 h APF and in the adult eye, Dpy and Pio were lost from the outer surface of the eye and only present in the pseudocone below the corneal lens in both wild-type and *dyl* mutant ommatidia (Fig. 5E, F, I). Dpy and Pio are thus in a position to provide outer and inner boundaries for the corneal lens as it is being secreted.

### Dpy and Pio are required for normal corneal lens shape

The Dpy protein, the largest isoforms of which have molecular weights close to 2.5 MDa, is known to promote membrane attachment to the cuticle in the embryonic trachea, the pupal wing, and pupal tendon cells, where it forms long filamentous structures ^39,40,43,49^. We hypothesized that Dpy might also contribute to attaching the corneal lens cuticle to the underlying cells. To test this idea, we generated clones homozygous for the lethal allele *dpy^lv^*^1^ ^36^ in the retina and examined them at pupal and adult stages. In adults, the overlying corneal lenses showed abnormal and irregular shapes (Fig. 6B, F, G), comparable to *dyl^Δ^*^42^ clones. However, *dpy* mutant ommatidia were not shorter on the apical-basal axis, consistent with the lack of significant apical constriction in *dpy* clones earlier in development (Fig. 6A, E). Clones mutant for the protein null allele *pio^V^*^132^ ^36^ (Fig. S6A) had no effect on apical constriction (Fig. 6C, Fig. S6A) and did not alter the external curvature of the corneal lens (Fig. 6D, F), but showed a deep and distorted inner corneal lens curvature (Fig. 6D, G). The protein null allele *tyn*^1^ ^62^ did not affect corneal lens shape (Fig. S6B-D). These results indicate that Dpy and Pio organized by Dyl have critical roles in determining corneal lens shape independent of apical cell size.

**Figure 6.**
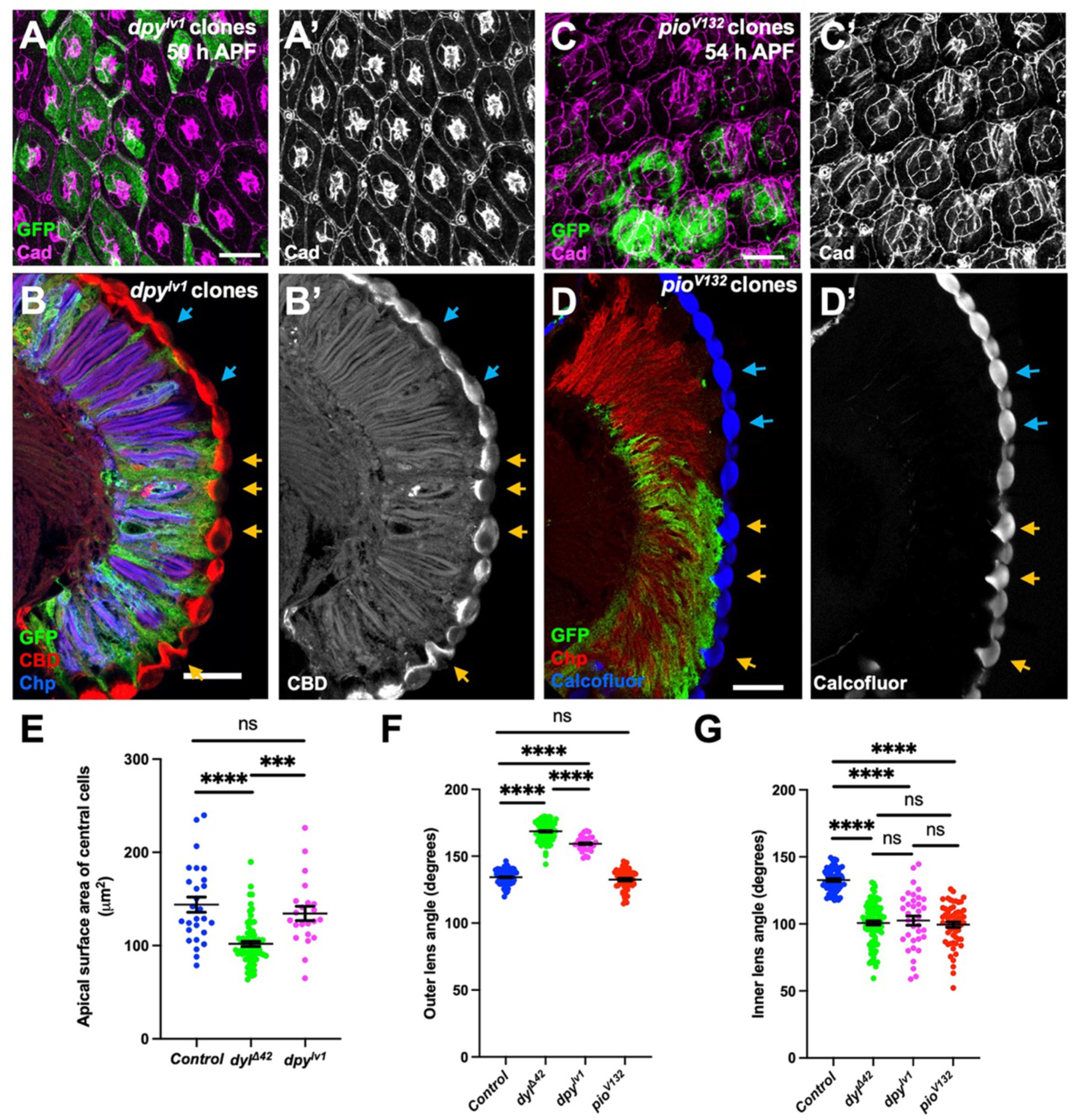
Loss of *dpy* or *pio* alters corneal lens shape. (**A, B**) 50 h APF retina (**A**) and adult eye section (**B**) containing *dpy^lv^*^1^ mutant clones labeled with GFP (green), stained for Ecad and Ncad (**A’**, magenta in **A**), Chp (blue in **B**) and CBD (**B’**, red in **B**). Although *dpy^lv^*^1^ mutant clones in the pupal retina show little apical constriction, corneal lenses in the adult eye have distorted shapes including changes in the inner and outer curvature (yellow arrows) in comparison to adjacent control corneal lenses (blue arrows). (**C, D**) 54 h APF retina (**C**) and adult eye section (**D**) containing *pio^V^*^132^ mutant clones labeled with GFP (green), stained for Ecad and Ncad (**C’**, magenta in **C**), Chp (red in **D**) and CBD (**B’**, blue in **B**). *pio* mutant clones do not show apical constriction at 54 h APF. In adults, *pio* mutant corneal lenses have normal outer curvature, but increased and distorted inner curvature (yellow arrows). (**E**) Graph showing the apical surface area of central cells in *dpy^lv^*^1^ mutant clones (n=22/10) compared to internal control wild-type ommatidia (n=27/10) and *dyl^11^*^42^ mutant clones (n=73/23) at 50 h APF. Although *dyl* mutant ommatidia are significantly constricted at this stage, *dpy* mutant ommatidia are not. (**F, G**) Graphs illustrating the outer angle (**F**) and inner angle (**G**) between neighboring corneal lenses in control, *dyl**^Δ^***^42^ mutant clones (data from Fig. 1F, G), *dpy^lv^*^1^ mutant clones (n=30/5) and *pio^V^*^132^ mutant clones (n=58/8) in the adult eye. Error bars show mean +/-SEM. ****p < 0.0001, ***p = 0.0004, ns p = 0.3965 (**E**), 0.1538 (**F**), 0.6419 (*dyl^11^*^42^ vs *dpy^lv^*^1^, **G**), 0.6217 (*dyl^11^*^42^ vs *pio^V^*^132^, **G**), 0.4525 (*dpy^lv^*^1^ vs *pio^V^*^132^, **G**), unpaired two-tailed t test with Welch’s correction. Scale bars: 10 µm (**A, C**), 20 µm (**B, D**).

### ZP-domain proteins act as a scaffold to retain chitin

A major component of the corneal lens is chitin, a polymer of N-acetylglucosamine. Chitin assembles into strong fibrillar structures that provide structural integrity ^63^. These chitin polymers are associated with numerous chitin-binding proteins, such as Gasp ^64^, which is highly expressed in the pupal retina ^20^. To examine whether the secretion or arrangement of chitin was affected in *dyl* mutants, we stained *dyl^Δ^*^42^ clones in the pupal retina with a fluorescently labeled CBD probe ^20^. We observed that at 50 h APF, chitin fibrils were located on the apical surface of wild-type ommatidia, but in *dyl^Δ^*^42^ clones chitin accumulation was sparse (Fig. S7A). At 54 h APF high levels of chitin were localized above wild-type ommatidia, but chitin was absent above *dyl* clones and did not appear to be stuck in the secretory pathway within the mutant cells (Fig. 7A, B). This phenotype could be rescued by expressing a *UAS-dyl* transgene in all retinal cells (Fig. S7H). Similarly, Gasp was not observed above *dyl^Δ^*^42^ clones at either 50 h (Fig. S7B) or 54 h APF (Fig. 7D). *dpy* mutant clones also showed a loss of chitin accumulation (Fig. 7E), despite normal Dyl localization (Fig. S7C), but *pio* was not required for chitin retention at this stage (Fig. S7D). We first detected chitin above *dyl* clones at 57 h APF, but at a lower level than in wild-type regions and without the appropriate curvature (Fig. 7C). All these data suggest that ZP-domain proteins are necessary for the timely assembly of chitin in the corneal lens. We hypothesize that an external scaffold that includes Dpy and Pio is crucial to retain the nascent chitin layer and mold it into a biconvex shape. Changes in the arrangement of these proteins may allow chitin to diffuse away at the stage when it would normally be deposited (Fig. 7H). This external scaffold is lost, perhaps through proteolytic degradation ^65,66^, later in development when the chitinous corneal lens becomes a strong, rigid structure (Fig. 5E, F, I). The delay in chitin assembly in *dyl* mutant clones may explain the defects in the surface structure of the adult corneal lens (Fig. 1A).

**Figure 7.**
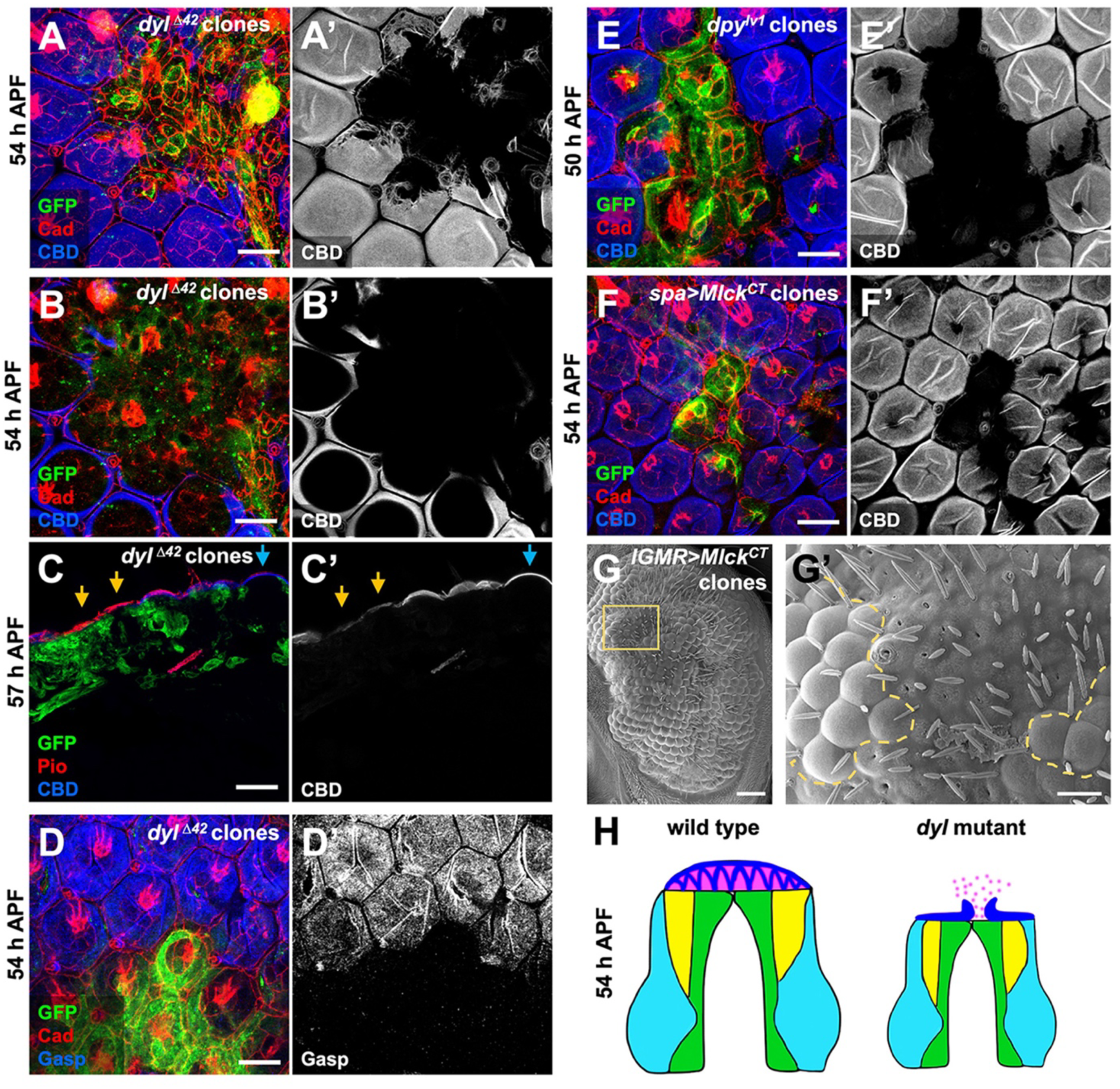
ZP-domain proteins and apical expansion are necessary for chitin accumulation. (**A-D**) Pupal retinas with *dyl^11^*^42^ clones marked with GFP (green), stained with CBD (**A’-C’**, blue in **A-C**) or anti-Gasp (**D’**, blue in **D**) and anti-Ecad and Ncad (red, **A, B, D**) or anti-Pio (red, **C**) at (**A**, **B, D**) 54 h APF and (**C**) a horizontal cryosection at 57 h APF. (**B**) is a more basal plane of the retina shown in (**A**). At 54 h APF chitin and Gasp are completely absent above *dyl* mutant cells. This decrease in accumulation is not due to a block in secretion, as no intracellular chitin is detected in the *dyl**^Δ^***^42^ mutant clones. Chitin begins to appear above the *dyl**^Δ^***^42^ mutant clones at 57 h APF, internal to a layer of Pio (yellow arrows in **C** and **C’**). (**E**) 50 h APF pupal retina with *dpy^lv^*^1^ mutant clones marked by GFP (green) and (**F**) 54 h APF retina with clones overexpressing *Mlck^CT^* with *spa-GAL4* marked by GFP (green) stained with anti-Ecad and Ncad (red) and CBD (**E’**, **F’**, blue in **E**, **F**). Loss of *dpy* and apical constriction induced by *Mlck^CT^* result in a similar loss of chitin. (**G**) Scanning electron micrograph of an eye containing clones expressing *UAS-Mlck^CT^* with *lGMR-GAL4*. The boxed area is enlarged in (**G’**), and the border of the clone is indicated with a dashed yellow line. The corneal lenses in *Mlck^CT^*-expressing ommatidia have flat external surfaces, but there are few gaps in the surface layer. Scale bars: 10 µm (**A-F**), 50 μm (**G**), 20 μm (**G’**). (**H**) Model depicting horizontal sections of wild type and *dyl* mutant ommatidia at 54 h APF. Chitin (pink) is retained by an external scaffold of proteins including Dpy and Pio (blue) in wild type ommatidia, but when the scaffold is disorganized in the absence of Dyl or Dpy, chitin can diffuse away.

Interestingly, we observed that chitin was also absent and Dpy was disorganized at 54 h APF above clones in which apical constriction of cone and primary pigment cells was induced by expressing activated Mlck (Fig. 7F, Fig. S7G). This suggests that the cell shape changes caused by loss of *dyl* may be sufficient to explain the altered ZP-domain scaffold and the lack of chitin retention. It is also possible that direct interactions with Dpy and Pio contribute to the effect of Dyl on their organization, as partially rescuing the constriction defect of *dyl* mutant clones by knocking down *Mlck* failed to rescue Pio organization or chitin accumulation (Fig. S7E, F). In addition, scanning electron micrographs confirmed the flat external surfaces of ommatidia expressing Mlck^CT^, but did not show the extensive surface gaps that were seen in *dyl* mutant clones (Fig. 7G). These results suggest that Dyl assembles a ZP-domain protein structure that prevents dispersal of the components of the developing corneal lens primarily by maintaining apical expansion of cone and primary pigment cells, but potentially also through an independent mechanism.

## Discussion

We show here that forming the precisely curved architecture of the *Drosophila* corneal lens from apical ECM requires the ZP-domain protein Dyl. Dyl acts transiently at a critical point in development to maintain the apical expansion of corneal lens-secreting cells and to assemble a scaffold containing ZP-domain proteins that acts as a convex outer boundary within which chitin and other corneal lens components are retained. Our results add to a growing body of evidence that the shapes of rigid structures composed of aECM rely on specific sites of attachment to the underlying cells that are mediated by ZP-domain proteins ^39,45,46,67^.

### Cell shape determines apical ECM shape

Our data show that the dramatic apical expansion of primary pigment cells in the pupal retina ^18,68^ is essential to build a foundation for corneal lens assembly. Reversing this expansion either by removing Dyl or by activating myosin contraction results in severe corneal lens defects. The apicobasal contraction that occurs in *dyl* mutant or *Mlck^CT^*-expressing ommatidia places the apical surfaces of cone and primary pigment cells in a more basal position than their wild-type neighbors. If secondary and tertiary pigment cells retain their attachments to the aECM, the ommatidial surface would take on a concave shape, potentially explaining the deeper inner curvature of the overlying corneal lenses (Fig. 2H). The smaller surface area of the central cells could also directly alter the shape of the corneal lens if this structure is assembled one layer at a time, like the tectorial membrane ^69^. Such a pattern of assembly would be consistent with labeling of the outer surface of the early corneal lens and the inner surface of the adult corneal lens by our chitin-binding probe (Fig. 5H, I), if it preferentially binds to newly deposited chitin. Constriction or folding of the apical cell surface has been shown to produce ridges of aECM in the *C. elegans* cuticle and the *Drosophila* trachea ^14,15^; our results suggest that apical expansion can also drive the morphogenesis of specific aECM structures.

The striking apical constriction of cone and primary pigment cells that we observed in *dyl* mutants implies that Dyl is required to maintain their expanded shape. Dyl is located on the apical surfaces of these cells, and it and other ZP-domain proteins have been shown to attach the plasma membrane to the apical ECM ^36,37,42,46,70,71^. These attachments can alter cell shape; for instance, apical expansion of cells in the pupal wing is dependent on the ZP-domain proteins Miniature and Dusky ^72^, NOAH-1 and NOAH-2 are required for the concerted cell shape changes that drive elongation of the *C. elegans* embryo ^73^, and Hensin is needed to convert flat β- to columnar α-intercalated cells in the kidney ^74^. The NOAH-1, NOAH-2 and FBN-1 ZP-domain proteins in the embryonic sheath resist deformation of the *C. elegans* epidermis by mechanical forces ^70,73^. We suggest that attachment to the aECM by Dyl enables cone and primary pigment cells to resist mechanical tension exerted by the actomyosin cytoskeleton. Active myosin and β_H_-spectrin accumulate in *dyl* mutant cells, and reducing the level of these components partially rescues apical constriction. However, these proteins appear disorganized and do not form foci like those seen in the wild-type cells. Similar foci are associated with changes in apical cell area driven by pulsatile constriction and ratcheting ^56–58^, but continuous constriction can occur with a diffuse apical actomyosin distribution, for instance in boundary larval epithelial cells ^53,75^. The loss of apical anchorage may cause *dyl* mutant cells to constrict in a continuous, passive manner that does not involve controlled ratcheting.

### ZP-domain proteins act as a scaffold for chitin retention

Our data also reveal a role for other ZP-domain proteins, including Dpy and Pio, in establishing an external scaffold that helps to shape the corneal lens. Although *dpy* mutant ommatidia do not show significant constriction, indicating that *dpy* acts downstream of or in parallel to the cell shape changes, they fail to accumulate chitin at the normal time and produce deformed corneal lenses. As chitin is not found trapped inside the mutant cells, it is likely that it diffuses away in the absence of the scaffold. Although Pio is not necessary to retain chitin at this stage, it appears to have an analogous role later in development, when it is present in the pseudocone, to maintain the normal morphology of chitin at the inner surface of the corneal lens (Fig. 6D). Dpy and Pio have a similar function in the trachea, where they provide an elastic structural element of the lumen that controls its shape and size and maintains the chitin matrix ^66,76,77^. Removal of this structure by proteolysis is necessary for subsequent gas filling of the airway ^65,66^, and it is possible that the external corneal lens scaffold is likewise proteolytically degraded in late pupal stages. Transient aECM structures containing ZP-domain proteins also shape many cuticular and tubular structures in *C. elegans* ^7^, and α-tectorin on the surface of supporting cells templates the assembly of collagen fibrils in each layer of the tectorial membrane ^69^.

We find that the arrangement of Dpy and Pio is dependent on Dyl; in wild-type ommatidia, they form linear structures, but this organization is not apparent in *dyl* mutants. We do not know the precise nature of these structures, nor what additional components they may contain. ZP-domain proteins are capable of forming heteropolymeric filaments ^31–33^, and another ZP protein, Qsm, is known to promote the secretion and remodeling of Dpy filaments that link tendon cells to the pupal cuticle, increasing their tensile strength ^40^. Since Dyl is only present in the retina for a short time, it may be required to initiate the assembly of a scaffold structure that then becomes self-sustaining. Alternatively, since apical constriction is sufficient to alter Dpy organization and delay chitin accumulation, the role of Dyl may be limited to maintaining apical expansion. Nevertheless, the requirement for *dyl* for normal chitin deposition in bristles and wing hairs ^47,48^ suggests that promoting chitin assembly, directly or indirectly, is one of its primary functions.

Other components of the corneal lens may also be affected by loss of *dyl*. Although chitin levels appear normal in sections of adult retinas, scanning electron micrographs show that the external surface of the corneal lens is incomplete. Induced apical constriction does not fully reproduce this loss of surface structure, suggesting that it reflects an independent function of Dyl. In the pupal wing, *dyl* belongs to a cluster of genes with peak expression at 42 h APF, when the outer envelope layer of the cuticle is being deposited, but it influences the structure of inner layers as well ^78^. Another gene expressed at this time point, *tyn*, is required for the barrier function of the embryonic envelope layer ^62^. It is not clear whether the corneal lens has a typical cuticular envelope, but its outermost layer forms nanostructures composed of the Retinin protein and waxes ^79^. Although *retinin* mRNA is not strongly expressed until very late in pupal development ^80^, its expression is initiated at mid-pupal stages when Dyl is present ^16^. It is possible that Dyl and the Dpy-Pio scaffold also control the retention of Retinin and other components of the outer layer.

In summary, we show here that the development of a biconvex corneal lens depends on both its upper and lower boundaries. The upper boundary, a convex shell of aECM that contains the ZP-domain proteins Dpy and Pio, encloses chitin and its associated proteins as they are secreted by cone and pigment cells. The shape of this shell may be defined by its attachment to the peripheral lattice cells and the pressure exerted on its center by secreted corneal lens components. The lower boundary is the apical plasma membrane of the cone and primary pigment cells. This membrane is flat until pseudocone secretion initiates late in pupal development, and the phenotype of *dyl* mutants suggests that it is maintained taut and expanded by attachment to the aECM. Loss of this expansion or of the pseudocone components Dpy or Pio gives the inner surface of the corneal lens a deeper curvature. It would be interesting to investigate whether mechanical forces exerted on the stromal extracellular matrix of the human cornea, perhaps through the ZP-domain protein ZP4 present in the underlying corneal endothelium ^81^, affect its shape and refractive power ^82,83^.

## STAR Methods

### Key resource table

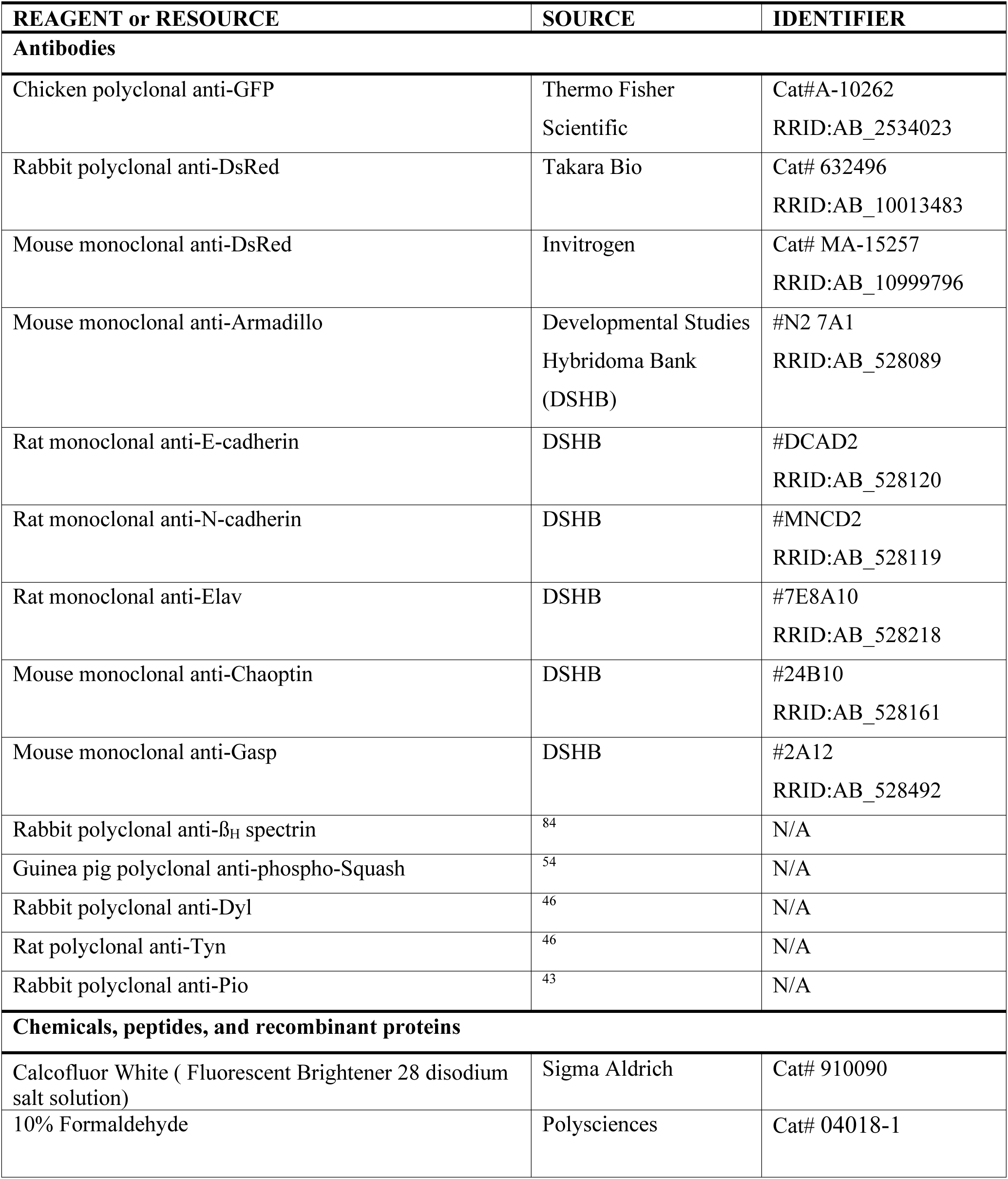
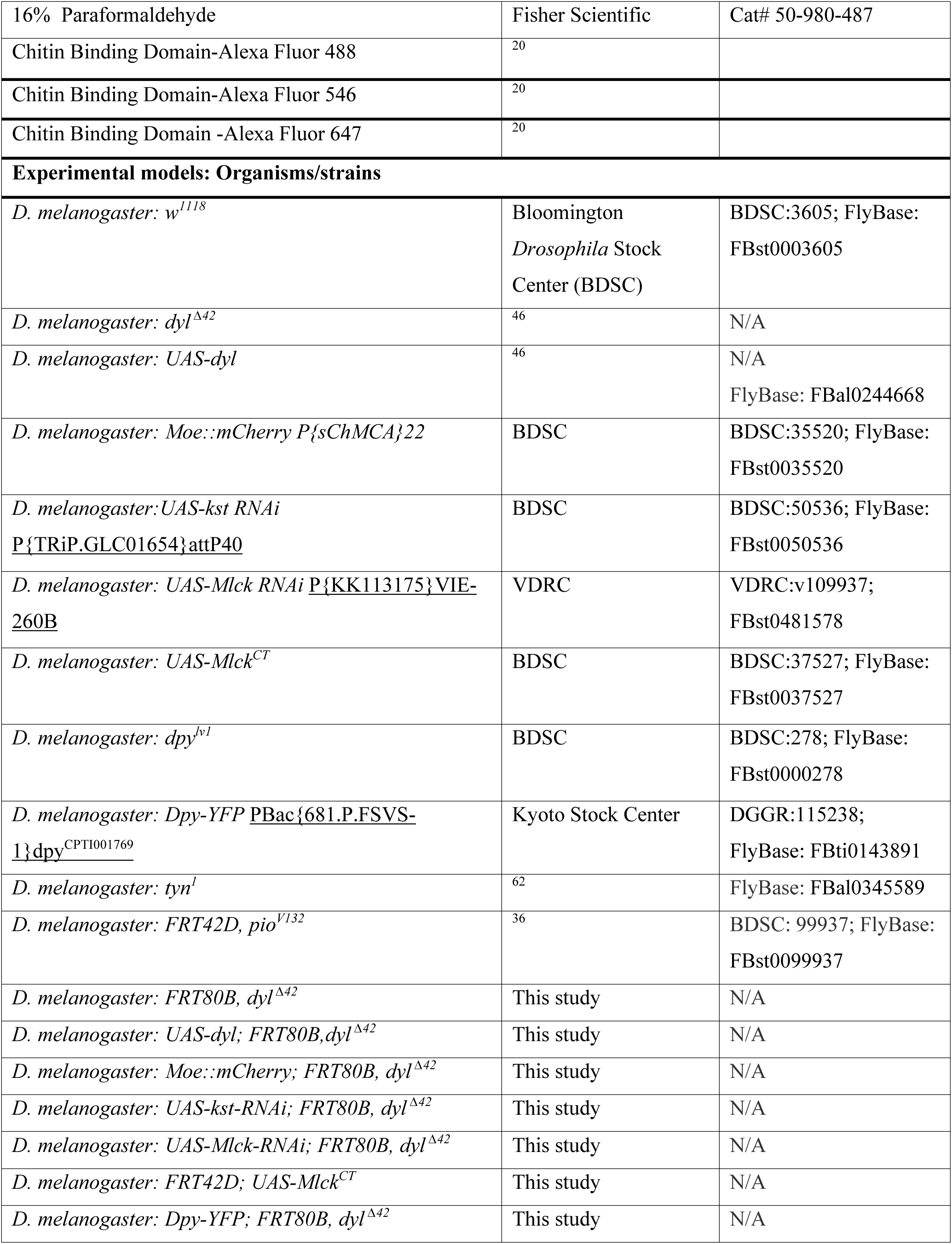
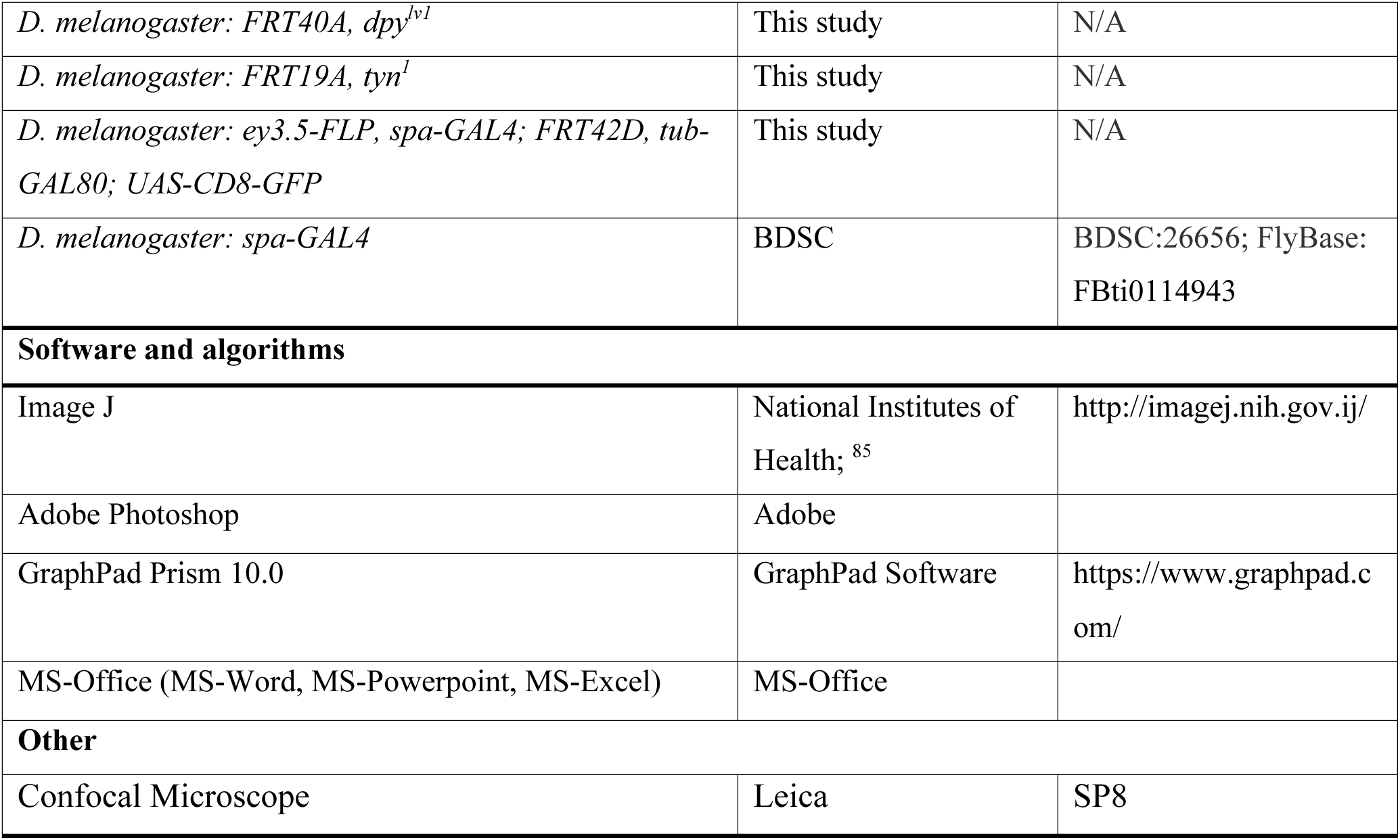

### Resource availability

#### Lead contact

Further information and requests for resources and reagents should be directed to and will be fulfilled by the lead contact, Jessica E. Treisman.

#### Materials availability

All flies and custom reagents created for this study are available upon request to the lead contact.

### Experimental model details

#### Fly stocks and genetics

*Drosophila melanogaster* strains were maintained on standard yeast-cornmeal-agar media and raised at 25°C. For analysis of the pupal retina, white prepupae (0 h APF) were collected with a soft wetted brush and cultured at 25°C till the appropriate developmental stage. Both sexes were used interchangeably for all the experiments, as no sex-specific differences were observed.

*Drosophila melanogaster* stocks used to generate *dyl ^Δ^*^42^ mutant clones were: (1) *UAS-CD8-GFP, ey3.5-FLP; lGMR-GAL4; FRT80B, tub-GAL80/TM6B*; (2) *FRT80B*, *dyl^Δ^*^42^*/TM6B*; (3) *Moe::mCherry; FRT80B*, *dyl ^Δ^*^42^*/SM6.TM6B; (4) Dpy-YFP; FRT80B*, *dyl ^Δ^*^42^*/SM6.TM6B*; (5) *eyFLP; lGMR-GAL4, UAS-myrTomato; FRT80B, tub-GAL80/TM6B;* (6) *UAS-dyl; FRT80B*, *dyl^Δ^*^42^*/TM6B;* (7) *UAS-kst RNAi; FRT80B*, *dyl ^Δ^*^42^*/SM6.TM6B* (8) *UAS-Mlck RNAi; FRT80B, dyl^Δ^*^42^*/SM6.TM6B*. Stocks used to generate *Mlck^CT^* overexpression clones were: (1) *UAS-CD8-GFP, ey3.5-FLP; lGMR-GAL4; FRT42D, tub-GAL80/TM6B*; (2) *ey3.5-FLP, spa-GAL4; FRT42D, tub-GAL80; UAS-CD8-GFP; (3) FRT42D; UAS-Mlck^CT^ /SM6.TM6B*. Stocks that were used to generate *dpy^lv^*^1^ mutant clones were: (1) *UAS-CD8-GFP, ey3.5-FLP; FRT40A, tub-GAL80; lGMR-GAL4 /TM6B*; (2) *FRT40A*, *dpy^lv^*^1^*/SM6.TM6B.* Stocks that were used to generate *pio^V^*^132^ mutant clones were: (1 *UAS-CD8-GFP, ey3.5-FLP; lGMR-GAL4; FRT42D, tub-GAL80/TM6B*; (2) *FRT42D*, *pio^V^*^132^*/ SM5.* Stocks that were used to generate *tyn*^1^ mutant clones were: (1) *ey3.5-FLP, FRT19A, tub-GAL80; lGMR-GAL4, UAS-CD8-GFP /SM6-TM6B*; (2) *FRT19A*, *tyn*^1^*/ FM7, Tb, Ubi-RFP*.

#### Immunohistochemistry

For cryosectioning, adult or pupal heads with the proboscis removed were glued onto glass rods using nail polish and fixed for 4 h in 4% formaldehyde in 0.2 M sodium phosphate buffer (pH 7.2) (PB) at 4°C. The heads were then incubated through an increasing gradient of sucrose in PB (5%, 10%, 25%, and 30% sucrose) for 20 min each, transferred to plastic molds containing OCT compound and frozen on dry ice. Cryosections of 12 µm were cut at −21°C, transferred onto positively charged slides and postfixed in 0.5% formaldehyde in PB at room temperature (RT) for 30 mins. The slides were then washed in PBS with 0.3% Triton X-100 (PBT) three times for 10 min each, blocked for 1 h at RT in 1% bovine serum albumin (BSA) in PBT and incubated in primary antibodies overnight at 4°C in 1% BSA in PBT. After three 20-minute washes in PBT, slides were incubated in secondary antibodies in 1% BSA in PBT for 2 h at RT and mounted in Fluoromount-G (Southern Biotech). A 1:5 dilution of Calcofluor White solution (25% in water; Sigma Aldrich, 910090) was included with the secondary antibodies where indicated.

Pupal retinas attached to the brain were dissected from staged pupae and collected in ice-cold PBS in a glass plate. These samples were fixed on ice in 4% formaldehyde in PBS for 30 min. The samples were washed three times for 10 min each in PBT and incubated overnight at 4°C in primary antibodies in 10% donkey serum in PBT. After three 20-min washes in PBT, the samples were incubated for 2 h in secondary antibodies in PBT/10% serum at RT, and washed again three times for 20 min in PBT. Finally, the retinas were separated from the brain and mounted in 80% glycerol in PBS.

The primary antibodies used were: mouse anti-Chp (1:50; Developmental Studies Hybridoma Bank (DSHB), 24B10), chicken anti-GFP (1:400; Thermo Fisher, A-10262), rat anti-Elav (1:100; DSHB, Rat-Elav-7E8A10), rat anti-Ecad (1:10, DSHB, DCAD2), mouse anti-Gasp (1:20, DSHB, 2A12), rabbit anti-Dyl (1:300) ^46^, rat anti-Tyn (1:100) ^46^, rabbit anti-Pio (1:300) ^43^, guinea pig anti-pSqh (1:1000) ^54^, and rabbit anti-ß_H_-spectrin (1:5000) ^84^. All antibodies were validated either using mutant or knockdown conditions as shown or by verifying that the staining pattern matched previously published descriptions. The secondary antibodies used were from either Jackson ImmunoResearch (Cy3 or Cy5 conjugates used at 1:200) or Invitrogen (Alexa488 conjugates used at 1:1000). Fluorescently labeled SNAP-CBD-probes (1:200) ^20^ were included with the secondary antibodies. Images were acquired on a Leica SP8 confocal microscope with a 63X oil immersion lens and processed using ImageJ and Adobe Photoshop.

#### Electron microscopy

For transmission electron microscopy, adult heads were cut in half and incubated in freshly made fixative containing 2.5% glutaraldehyde, 2% paraformaldehyde, and 0.05% Triton X-100 in 0.1 M sodium cacodylate buffer pH 7.2 (CB) on a rotator for at least 4 h until all heads had sunk to the bottom of the tube, and then in the same fixative without Triton X-100 overnight at 4°C on a rotator. After washing three times for 10 min in CB, the heads were post-fixed in 1% OsO_4_ in CB for 1.5 h, washed three times for 10 min in water, dehydrated in an ethanol series (30%, 50%, 70%, 85%, 95%, 100%), rinsed twice with propylene oxide and embedded in EMbed812 epoxy resin (Electron Microscopy Sciences). 70 nm ultrathin sections were cut and mounted on formvar-coated slot grids and stained with uranyl acetate and lead citrate ^86^. Electron microscopy imaging was performed on a Talos120C transmission electron microscope (Thermo Fisher Scientific) and recorded using a Gatan (4k×4k) OneView Camera with Digital Micrograph software (Gatan).

For scanning electron microscopy, whole flies were fixed for 2 h at room temperature and then overnight at 4°C in 1% glutaraldehyde/1% formaldehyde/0.2% Triton X-100/0.1 M sodium cacodylate pH 7.2. After three 10 min washes in PBS, the flies were postfixed in 1% OsO4 in CB for 1 h and rinsed three times for 10 min in water. They were dehydrated in an ethanol series (30%, 50%, 70%, 85%, 95%, 3X 100%) and stored in 100% ethanol overnight. The eyes were critical point dried using a Tousimis Autosamdri®-931 critical point dryer (Tousimis, Rockville, MD), mounted on SEM stabs covered with double sided electron-conductive tape, coated with gold/palladium by a Safematic CCU-010 SEM coating system (Rave Scientific, Somerset, NJ), and imaged on a Zeiss Gemini300 FESEM (Carl Zeiss Microscopy, Oberkochen, Germany) using a secondary electron detector (SE_2_) at 5 kV with working distance (WD) between 15.2 mm and 19.4 mm.

#### Quantification and statistical analysis

The outer and inner angles between adjacent corneal lenses were measured according to the schematic in Fig. 1E, using the angle tool in ImageJ. To measure the apical surface areas of ommatidia and central cells, ROI were drawn and measured in ImageJ using the freehand selection tool in Z-projections that included the apical surface of the retina. To measure ommatidial height, freehand straight lines were drawn from the corneal lens surface to the base of the rhabdomere using the line tool in ImageJ and measured. Values were plotted in GraphPad Prism v10. Significance was calculated using Welch’s two-tailed unpaired *t*-tests. Sample numbers and definitions of error bars are given in the figure legends. Sample sizes for quantifications were not predefined and no samples were excluded.

## Data and code availability

This study did not generate or analyze any datasets/codes. Any additional information required to reanalyze the data reported in this paper is available from the Lead Contact upon request.

## Acknowledgments

We thank Markus Affolter, Shigeo Hayashi, François Payre, Robert Ward, Christian Klämbt, the Bloomington *Drosophila* stock center, the Vienna *Drosophila* resource center, the Kyoto stock center, and the Developmental Studies Hybridoma Bank for fly stocks and reagents. Information available on FlyBase was invaluable for this work. We thank NYULH DART Microscopy Laboratory members Alice F.-X. Liang, Joseph Sall, Jason Liang and Chris Petzold for consultation and assistance with electron microscopy. This core is partially funded by NYU Cancer Center Support Grant NIH/NCI P30CA016087, and the Gemini300 FESEM was supported by NIH S10 OD019974. We thank Dhaval Gandhi, Sharukh Khan and Genie Jang for technical assistance. The manuscript was improved by the critical comments of Gira Bhabha, Maria Bustillo, Holger Knaut, Sudershana Nair, and Hongsu Wang. This work was funded by the National Institutes of Health (grant R01EY032896 to J.E.T.).

## Author contributions

Conceptualization, N.G. and J.E.T.; investigation, N.G.; data curation, N.G.; formal analysis, N.G. and J.E.T.; writing-original draft, N.G.; writing – review and editing, J.E.T.; funding acquisition, J.E.T.; supervision, J.E.T.

