## Supplementary Figures for "Apical cell expansion maintained by Dusky-like establishes a scaffold for corneal lens morphogenesis"

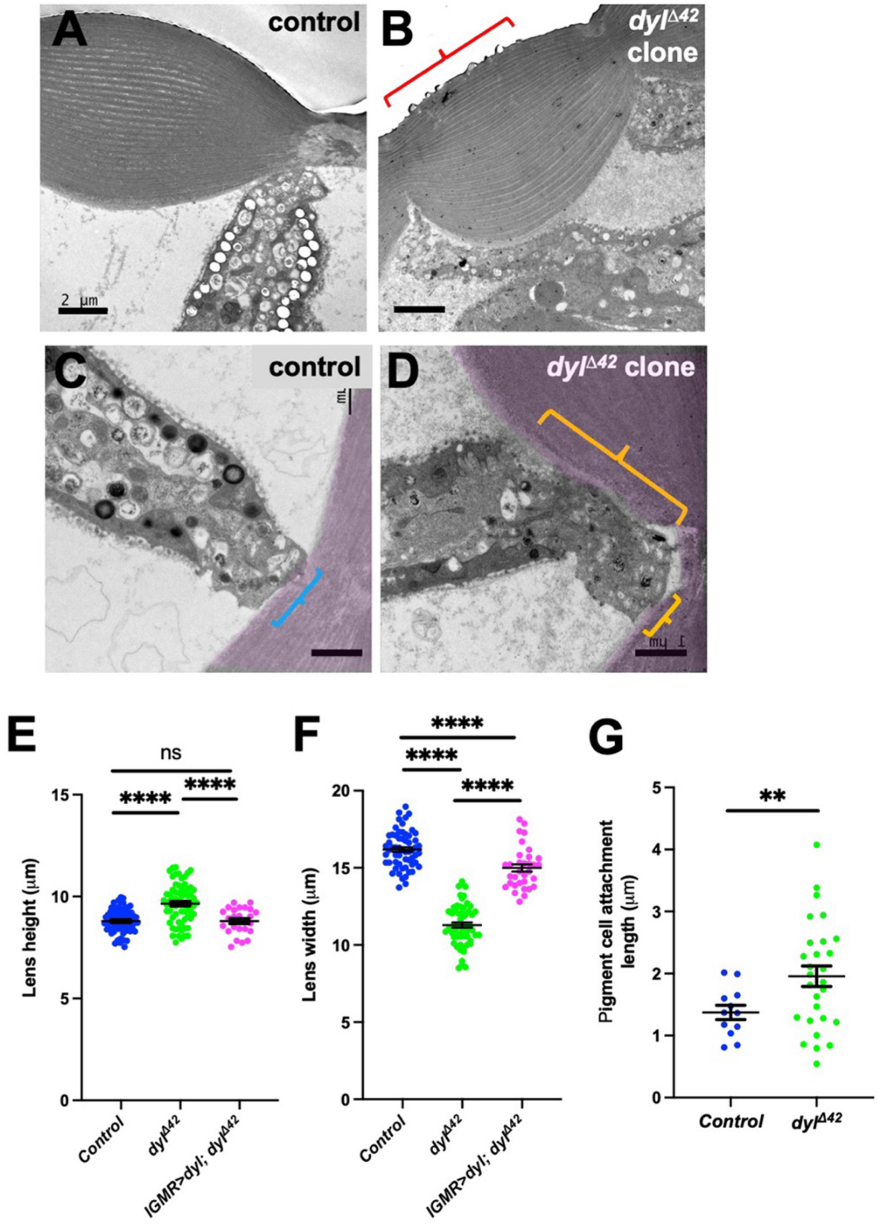


**Figure S1. Changes in corneal lens morphology in *dyl* mutants.** (**A-D**) transmission electron micrographs of wild-type (**A, C**) and *dyl*$^{\Delta 42}$ mutant (**B, D**) ommatidia in adult eyes. (**B**) shows defects and protrusions on the external surface of the *dyl* mutant corneal lens (bracket). (**C, D**) show sites of contact (brackets) between the pigment cells and the corneal lens (pseudocolored in magenta). Scale bars: 2 µm (**A, B**), 1 µm (**C, D**). (**E, F**) graphs showing corneal lens height (**E**) and corneal lens width (**F**) in wild type control regions of mosaic adult eye sections (n=73/11), *dyl*$^{\Delta 42}$ mutant clones (n=68/11), and *dyl*$^{\Delta 42}$ mutant clones expressing *UAS-dyl* with *lGMR-GAL4* (n=37/6). (**G**) Graph showing the length of pigment cell-corneal lens contacts in electron micrographs of control retinas (n=12/2) and *dyl*$^{\Delta 42}$ mutant clones (n=28/1). For all graphs, error bars show mean ± SEM. ****$p < 0.0001$, **p=0.0061, ns p=0.993, unpaired two-tailed t test with Welch's correction.

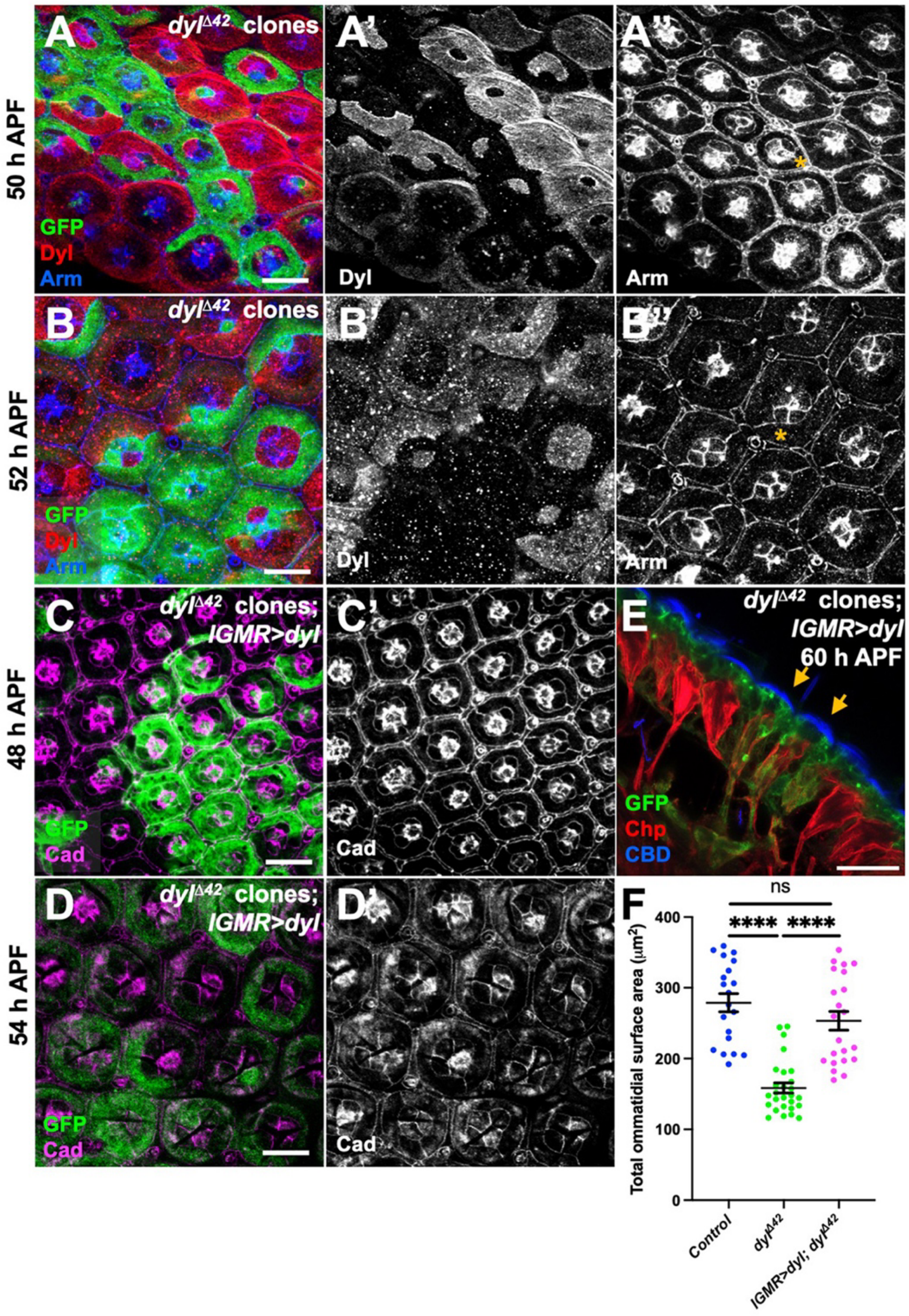


**Figure S2. Progression of the *dyl* mutant phenotype and rescue by UAS-*dyl*.** (**A, B**) Retinas containing $dyl^{\Delta42}$ mutant clones marked with GFP (green), stained with anti-Dyl (**A'**, **B'**, red in **A**, **B**) and anti-Arm (**A''**, **B''**, blue in **A**, **B**). (**A**) 50 h APF; (**B**) 52 h APF. Dyl staining is absent from *dyl* mutant clones and is strongest at 50 h APF. Apical constriction of some *dyl* mutant central cells is marked with yellow asterisks. (**C, D**) Pupal retinas in which $dyl^{\Delta42}$ mutant clones expressing *UAS-dyl* with *lGMR-GAL4* are marked with GFP (green), stained with anti-E-cadherin (Ecad) and anti-N-cadherin (Ncad) to mark apical cell junctions (**C'**, **D'**, magenta in **C**, **D**). (**C**) 48 h APF; (**D**) 54 h APF. (**E**) Horizontal cryosection of a 60 h APF retina in which $dyl^{\Delta42}$ mutant clones expressing *UAS-dyl* with *lGMR-GAL4* are marked with GFP (green, yellow arrows). The corneal lens is labeled with CBD (blue) and photoreceptors with anti-Chp (red).

Expression of wild-type Dyl rescues the apical constriction and apical-basal contraction. Scale bars: 10 µm (**A-D**), 20 µm (**E**). (**F**) A graph depicting the total ommatidial apical surface area in control (n=20/7), *dyl*$^{\Delta 42}$ mutant clones (n=27/7), and *dyl*$^{\Delta 42}$ mutant clones rescued with *lGMR-GAL4* driving *UAS-dyl* at 54 h APF (n=22/8). *UAS-dyl* fully rescues apical constriction in *dyl*$^{\Delta 42}$ mutant clones. Error bars show mean +/-SEM. ****$p < 0.0001$, ns $p = 0.1744$, unpaired two-tailed t test with Welch's correction.

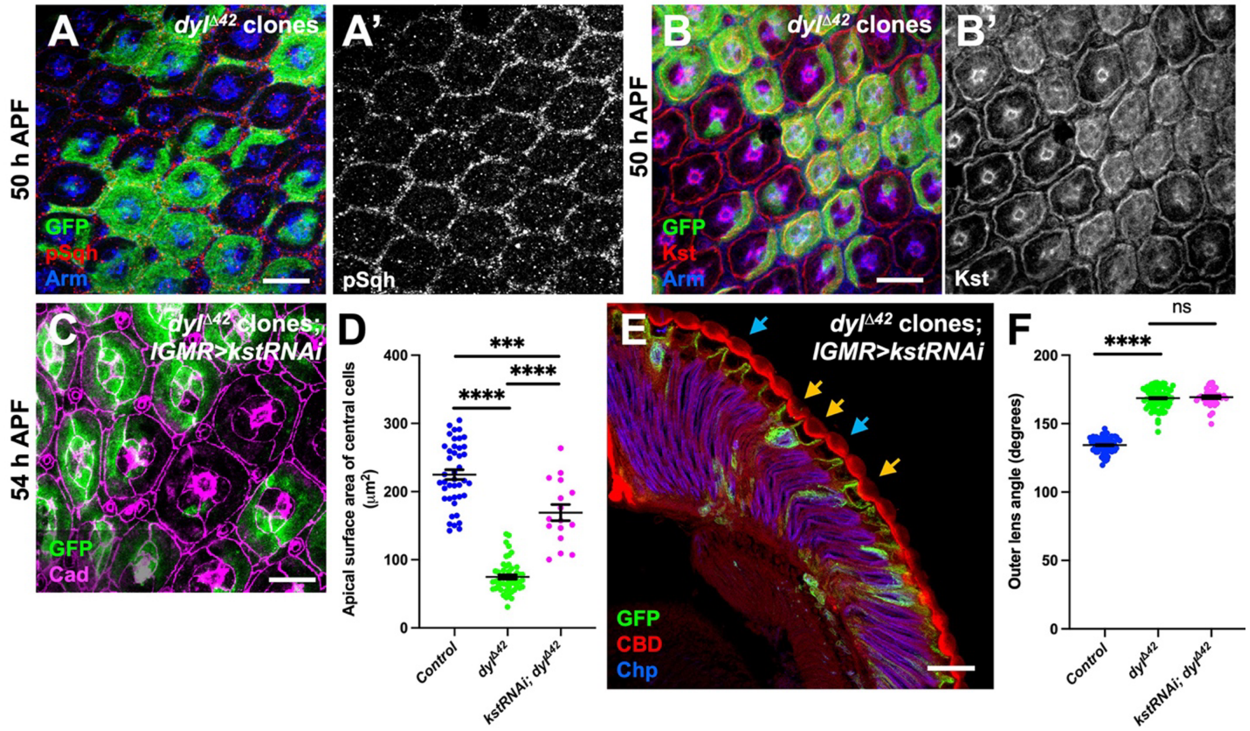


**Figure S3. Knocking down *kst* in *dyl* mutant clones does not rescue corneal lens curvature.** (**A, B**) 50 h APF retinas with *dyl*$^{\Delta 42}$ mutant clones marked with GFP (green), stained with anti-Arm (blue), anti-pSqh (**A'**, red in **A**) or anti-$\beta_H$-spectrin (**B'**, red in **B**). *dyl* mutant clones accumulate $\beta_H$-spectrin at this stage, but do not show a change in pSqh. (**C**) 54 h APF retina with *dyl*$^{\Delta 42}$ mutant clones expressing *UAS-kst RNAi* with *lGMR-GAL4*, marked with GFP (green). Apical cell outlines are stained with anti-Ecad and anti-Ncad (magenta). (**D**) Graph showing the apical surface area of central cells in internal wild type control ommatidia, *dyl*$^{\Delta 42}$ mutant clones (data from **Fig. 2D**), and *dyl*$^{\Delta 42}$ mutant clones expressing *kst RNAi* (n=16/7) at 54 h APF. (**E**) Cryosection of an adult eye expressing *kst* RNAi in *dyl* mutant clones marked with GFP (green), stained with anti-Chp (blue) and CBD (red). *kst* knockdown fails to rescue the corneal lens shape (yellow arrows) compared to wild-type corneal lenses (blue arrows). Scale bars: 10 µm (**A-C**), 20 µm (**E**). (**F**) Graph showing the outer angle between adjacent corneal lenses in control, *dyl*$^{\Delta 42}$ mutant clones (data from **Fig. 1F**), and *kst RNAi* in *dyl*$^{\Delta 42}$ mutant clones (n=35/6). Although knocking down *kst* in *dyl*$^{\Delta 42}$ mutant clones significantly rescues the apical surface area of central cells, it is not sufficient to restore normal corneal lens curvature. Error bars show mean +/-SEM. ****$p < 0.0001$, ***$p = 0.0004$, ns $p = 0.4159$, unpaired two-tailed t test with Welch's correction.

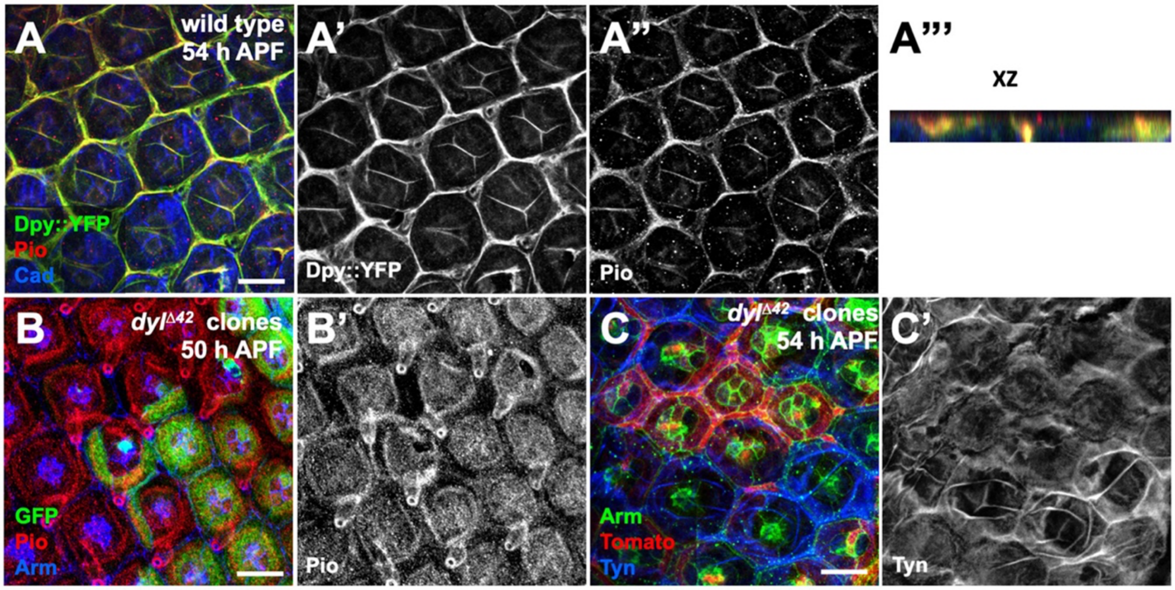


**Figure S5. Multiple ZP-domain proteins form an organized structure at the apical surface of the retina.** (**A**) A 54 h APF pupal retina showing colocalization of Dpy-YFP (**A', **green in **A, A'''**) and Pio (**A''**, red in **A, A'''**). Ecad and Ncad (blue) mark the apical junctions. An orthogonal section (**A'''**, apical up) shows that these proteins are in the same plane, apical to cadherin. (**B, C**) Pupal retinas containing *dyl*$^{\Delta 42}$ clones marked with GFP (green, **B**) or Tomato (red, **C**), stained with anti-Arm (blue in **B**, green in **C**), anti-Pio (**B'**, red in **B**), or anti-Tyn (**C'**, blue in **C**). (**B**) 50 h APF; (**C**) 54 h APF. Pio organization is not affected in *dyl*$^{\Delta 42}$ mutant clones at 50 h APF; Tyn shows a filamentous organization in wild-type but not *dyl* mutant ommatidia at 54 h APF. Scale bars: 10 µm.

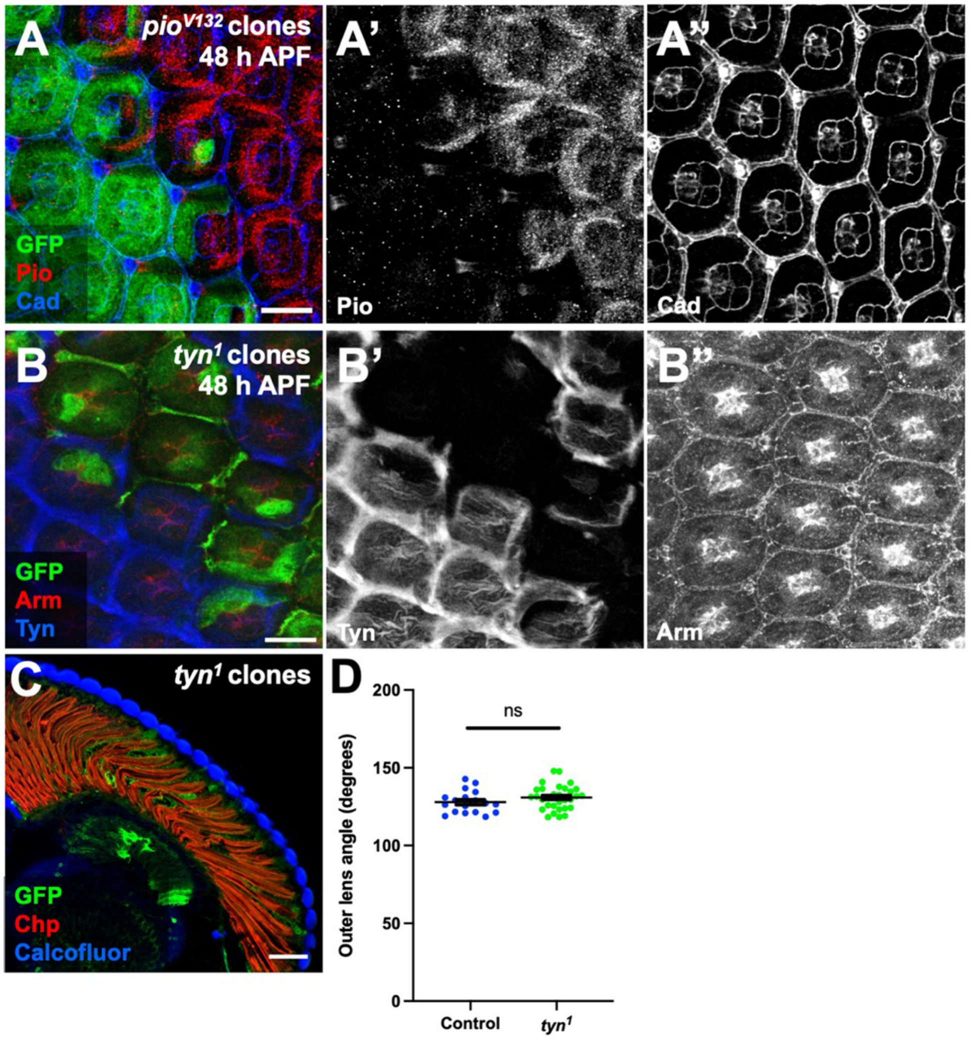


**Figure S6. Loss of *tyn* does not alter corneal lens shape.** (**A, B**) *pio*$^{V132}$ mutant clones (**A**) or *tyn*$^{1}$ mutant clones (**B**) marked with GFP (green) in 48 h APF pupal retinas labeled with anti-Ecad and anti-Ncad (**A'', blue in A)** or anti-Arm **(B''**, red in **B**) and anti-Pio (**A'**, red in **A**) or anti-Tyn (**B'**, blue in **B**). Both alleles are protein nulls. (**C**) Horizontal cryosection of an adult eye with *tyn*$^{1}$ mutant clones marked with GFP (green), labeled with anti-Chp (red) and Calcofluor White (blue). (**D**) A graph illustrating the outer lens angle between neighboring corneal lenses in internal wild type control (n=18/6) and *tyn*$^{1}$ mutant clones (n=31/6) in the adult eye. Error bars show mean +/-SEM. ns p = 0.185. Scale bars: 10 µm (**A, B**), 20 µm (**C**).

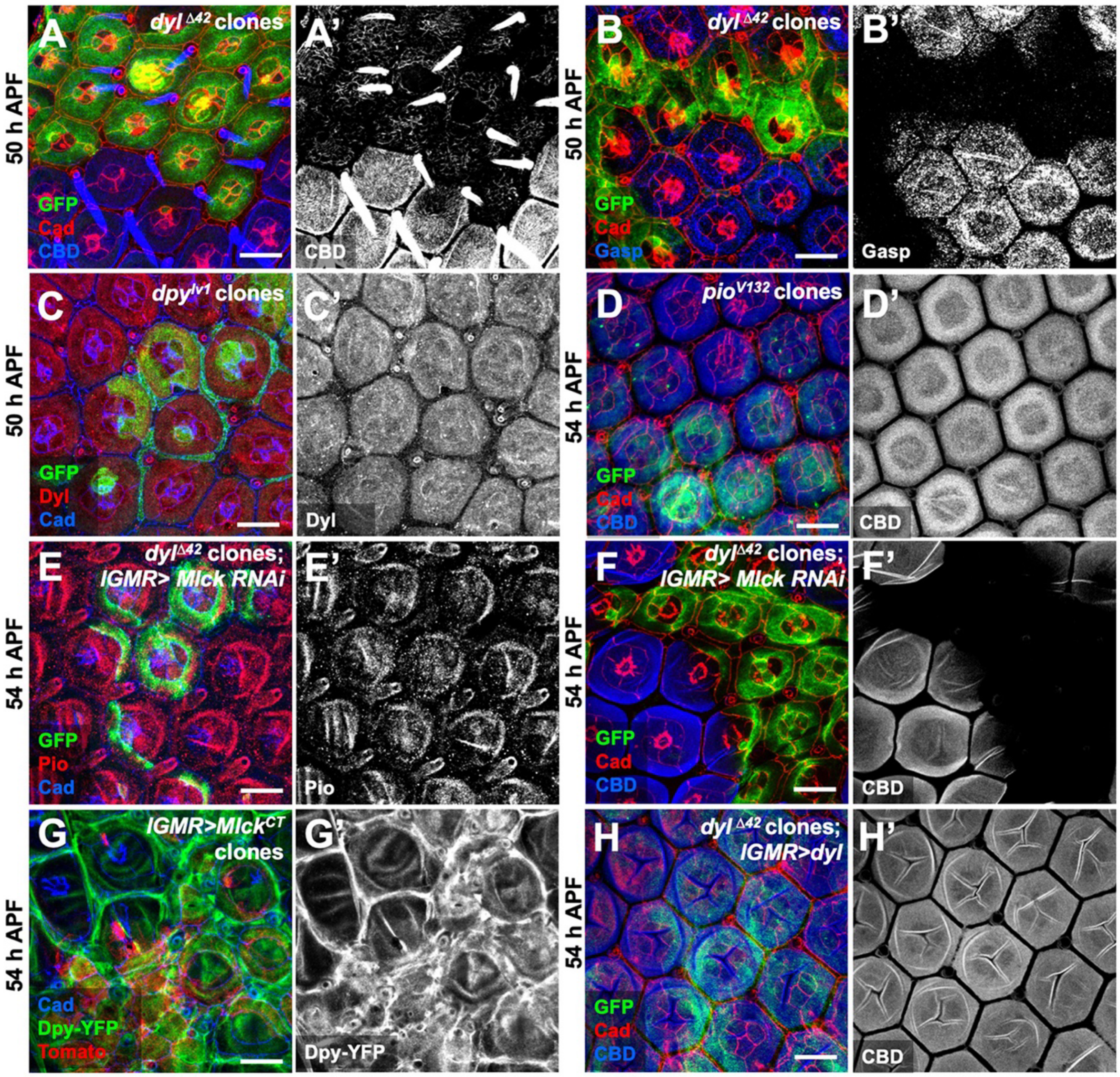


**Figure S7. Apical constriction acts on chitin through ZP-domain proteins.** (**A, B**) 50 h APF pupal retinas with *dyl*$^{\Delta 42}$ clones marked with GFP (green), stained with CBD (**A'**, blue in **A**), anti-Gasp (**B'**, blue in **B**), and anti-Ecad and Ncad (red). At 50 h APF there is a small amount of chitin above the *dyl*$^{\Delta 42}$ mutant clones compared to the more robust chitin accumulation above wild-type ommatidia, and Gasp accumulation in *dyl* mutant clones is already defective. CBD also labels mechanosensory bristles. (**C**) 50 h APF pupal retina with *dpy*$^{lv1}$ mutant clones marked with GFP (green), stained with anti-Dyl (**C'**, red in **C**) and anti-Ecad and Ncad (blue). Dyl localization is not affected in *dpy* clones. (**D**) 54 h APF pupal retina with *pio*$^{V132}$ mutant clones marked with GFP (green), stained with CBD (**D'**, blue in **D**) and anti-Ecad and Ncad (red). Loss of *pio* does not affect chitin accumulation at this stage. (**E, F**) *dyl*$^{\Delta 42}$ mutant clones expressing *UAS-Mlck RNAi* with *lGMR-GAL4*, marked by GFP (green) and stained with anti-Pio (**E'**, red in **E**), CBD (**F'**, blue in **F**) and anti-Ecad and Ncad (blue in **E**, red in **F**). Reduction of apical constriction in *dyl*$^{\Delta 42}$ mutant clones by knocking down *Mlck* does not rescue Pio organization or loss of chitin. (**G**) 54 h APF retina with clones expressing *UAS-Mlck*$^{CT}$ with *lGMR-GAL4* marked with Tomato (red), stained for Dpy-YFP (**G'**, green in **G**) and anti-Ecad and anti-Ncad (blue). Apical constriction is sufficient to alter Dpy organization. (**H**) 54 h APF retina with *dyl*$^{\Delta 42}$ mutant clones expressing *UAS-dyl* with *lGMR-GAL4* marked with GFP (green) stained with CBD (**H'**, blue in **H**) and Ecad and Ncad (red). Wild type *dyl* rescues chitin accumulation in *dyl*$^{\Delta 42}$ mutant ommatidia. Scale bars: 10 µm.
